# Hybrid-metagenomics reveal off-target effects of albendazole, ivermectin-albendazole and moxidectin-albendazole on the human gut bacteria

**DOI:** 10.1101/2025.06.27.660518

**Authors:** Julian Dommann, Viviane P. Sprecher, Christian Beisel, Daniel Ballmer, Eveline Hürlimann, Jean T. Coulibaly, Jennifer Keiser, Pierre H. H. Schneeberger

## Abstract

The human whipworm parasite, *Trichuris trichiura,* poses a critical public health problem, affecting over 400 million people globally and responding poorly to the standard-of-care - benzimidazole chemotherapy. Recent efforts aimed at developing improved treatment options, the two macrolide-benzimidazole combination therapies ivermectin-albendazole and moxidectin-albendazole. Recent reports suggest that ivermectin and moxidectin possess antibacterial properties *in vitro*, hence there is a need to comprehensively characterize their off-target effects on the gut microbiome. In the framework of a randomized-controlled trial we collected stool samples of 204 *T. trichiura*-infected individuals in Côte d’Ivoire receiving albendazole (400mg), albendazole-ivermectin (400mg/200µg/kg) and albendazole-moxidectin (400mg/8mg). Using a state-of-the-art hybrid sequencing approach that combines Illumina short read and Oxford Nanopore long read shotgun sequencing, we recovered over 800 high-quality metagenome-assembled genomes. Our analyses reveal that albendazole and albendazole-moxidectin induce taxonomic shifts in the gut microbiota with only mild functional consequences. In contrast, in individuals receiving higher quantities of albendazole-ivermectin (ivermectin > 12mg), based on their weight, we observed profoundly modulated taxonomic composition and microbial function, while the resistome was largely spared. These findings robustly confirm ivermectin’s antibacterial properties in the human gut, extending beyond previously reported *in vitro* effects. Given that ivermectin is a cornerstone in parasitic disease control, it is crucial to evaluate its broader impacts and considering more targeted, evidence-based treatment strategies.

## Introduction

Soil-transmitted helminthiasis, one of 21 neglected tropical diseases, affects over 1.5 billion people worldwide^1,2^. People living in resource-limited settings in the southern hemisphere are disproportionally affected. Noteworthy, soil-transmitted helminthiases encompasses the diseases caused by one or more species of gut-dwelling, parasitic nematodes. The soil-transmitted helminths (STHs) include the roundworm *Ascaris lumbricoides,* the threadworm *Strongyloides stercoralis*, the whipworm *Trichuris trichiura*, and the two hookworms *Ancylostoma duodenale* and *Necator americanus*. Depending on the parasite species and severity of infection, symptoms differ. A heavy infection may include symptoms such as intestinal obstructions (*A. lumbricoides*), rectal prolapse (*T. trichiura*) and severe anemia (hookworms)^3,4^. As all STHs require a part of their development period in soil and transmission mainly relies on contact with larvae-infested soil or foods, long-term disease prevention focuses on improving hygiene and sanitation^5^. The short-term remedy is based on mass drug administration (MDA) with albendazole (ALB) and mebendazole (MBZ) without prior diagnosis. In the case of soil-transmitted helminthiases, MDA focuses primarily on children and may take place annually or biannually. Importantly, species-level parasite epidemiology is heterogeneous, and drug efficacies against the varying parasites differ ^6,7^. For instance, single doses of ALB and MBZ are not sufficient to effectively reduce the burden and prevalence of *T. trichiura* infections^8^.

Like ALB and MBZ, ivermectin (IVM) is used in MDA programs to reduce the burden of lymphatic filariasis and onchocerciasis, two additional neglected tropical diseases^9,10^. IVM has recently been recommended as a partner-drug of ALB, potentially finding its way into STH MDA programs to broaden treatment efficacy against *T. trichiura*^11^. Moxidectin (MOX) is registered for onchocerciasis^12^ and might be used in combination with ALB for STH infections in the future^13–15^.

Structurally, both IVM and MOX, are macrocyclic lactones and therefore closely resemble the group of macrolide antibiotics. Owing to this fact, but also due to their shared habitat with the parasite in the host’s gastrointestinal tract, gut bacteria represent a tangible secondary target for IVM and MOX. In contrast to the parasites, the majority of gut bacteria make up an indispensable part of our gastrointestinal tract aiding in digestion, protection against pathogens and immune responses^16,17^. Disruption of the gut microbiome homeostasis could impede these essential functions. Given the harbored functional potential in the gut microbiome^18,19^ and the large-scale, continuous use of IVM - and at a later stage possibly MOX - in control programs, off-target activity may trigger bacterial drug resistance. While resistance to IVM in livestock helminths has been documented, no resistance has been detected in human nematodes or human gut bacteria^20,21^. However, recent studies demonstrated the wide-ranged antibacterial properties of the two anthelminthics and the phenotypical adaptation of bacteria to anthelminthics and multiple antibiotics after repeated anthelminthic exposure *in vitro*^22,23^.

Furthermore, analyses of the bacterial resistome after the trial “Macrolides Oraux pour Réduire les Décès avec un Oeil sur la Résistance I” (MORDOR I) demonstrated that biannual administration of the macrolide azithromycin for two years to reduce childhood mortality, increased the proportion of macrolide resistance genes in the gut microbiome^24,25^. Moreover, the proportion of macrolide resistance genes in the nasopharyngeal microbiome was elevated, demonstrating the role of the gut microbiome as a reservoir for antimicrobial resistance genes (ARGs) and emphasizing its key role in spreading ARGs^26^.

In light of potential future use of IVM and MOX against STH infections and the corresponding increase in administration, we aimed to identify the off-target effects of IVM and MOX on the bacterial gut microbiome characterized from stool samples collected in a randomized-controlled phase 3 trial conducted in Côte d’Ivoire in 2021^27^. In short, we generated pooled high-quality hybrid metagenomes for 204 participants infected with *T. trichiura,* spread across three treatment arms (ALB, IVM-ALB and MOX-ALB) and two timepoints (before treatment, BL; at follow-up after treatment, FU). In the subsequent analysis we compared, i) species-level taxonomic composition, ii) functional composition, as well as iii) the resistome between BL and FU and across treatment arms.

## Methods and Materials

### Sample origin

Stool samples were gathered as part of a randomized-controlled trial (NCT04726969; https://clinicaltrials.gov/study/NCT04726969) aimed at evaluating the efficacy and safety of MOX-ALB compared to IVM-ALB and ALB against *T. trichiura* in Côte d’Ivoire in 2021^27^ (Figure 1). Individuals interested in participating in the trial were invited to complete the process of informed consent; thereafter, individuals were assessed for study eligibility during screening procedures. Overall, 210 participants were randomly assigned to three treatment arms, namely ALB (n = 67), IVM-ALB (n = 71) and MOX-ALB (n = 72). For each participant, a small aliquot of stool (∼1g) was transferred to a sterile 2 mL cryotube, both before the treatment (BL) and 2-3 weeks after (FU). The collected aliquots were immediately frozen and kept at −20°C. Upon completion of the trial, these aliquots were shipped to Swiss TPH (Allschwil, Switzerland) on dry ice and kept at −80°C until DNA extraction. Only fully anonymized samples and isolates were used throughout the study and no participant-specific information was available. The sequencing analysis was limited to 204 paired samples, i.e. ALB (n = 67), IVM-ALB (n = 69) and MOX-ALB (n = 68), for which BL and FU samples were available and sequencing libraries could be produced. Since IVM treatment is weight-based (200µg/kg), we further stratified the IVM-ALB arm based on the weight of the subject. According to Maier *et al.*, oral administration of 11.4mg of IVM, which corresponds to an average amount of IVM following administration of 200µg/kg used for STH infections, result in an estimated intestinal IVM concentration of 4.6μM ^22^. Moreover, previously published *in vitro* studies have demonstrated that a diverse range of gut-bacterial isolates exhibit pronounced sensitivity to IVM at concentrations ≥ 5μM^23^. Therefore, we employed a weight threshold of 60kg, corresponding to 12mg of IVM, to separate the initial IVM-ALB cohort into a low-concentration group (IVM-ALB-LOW, n = 35) and a high-concentration group (IVM-ALB-HIGH, n = 34). These group will be referred to as arms, even though they did not represent a treatment arm of the trial per se.

**Figure 1:**
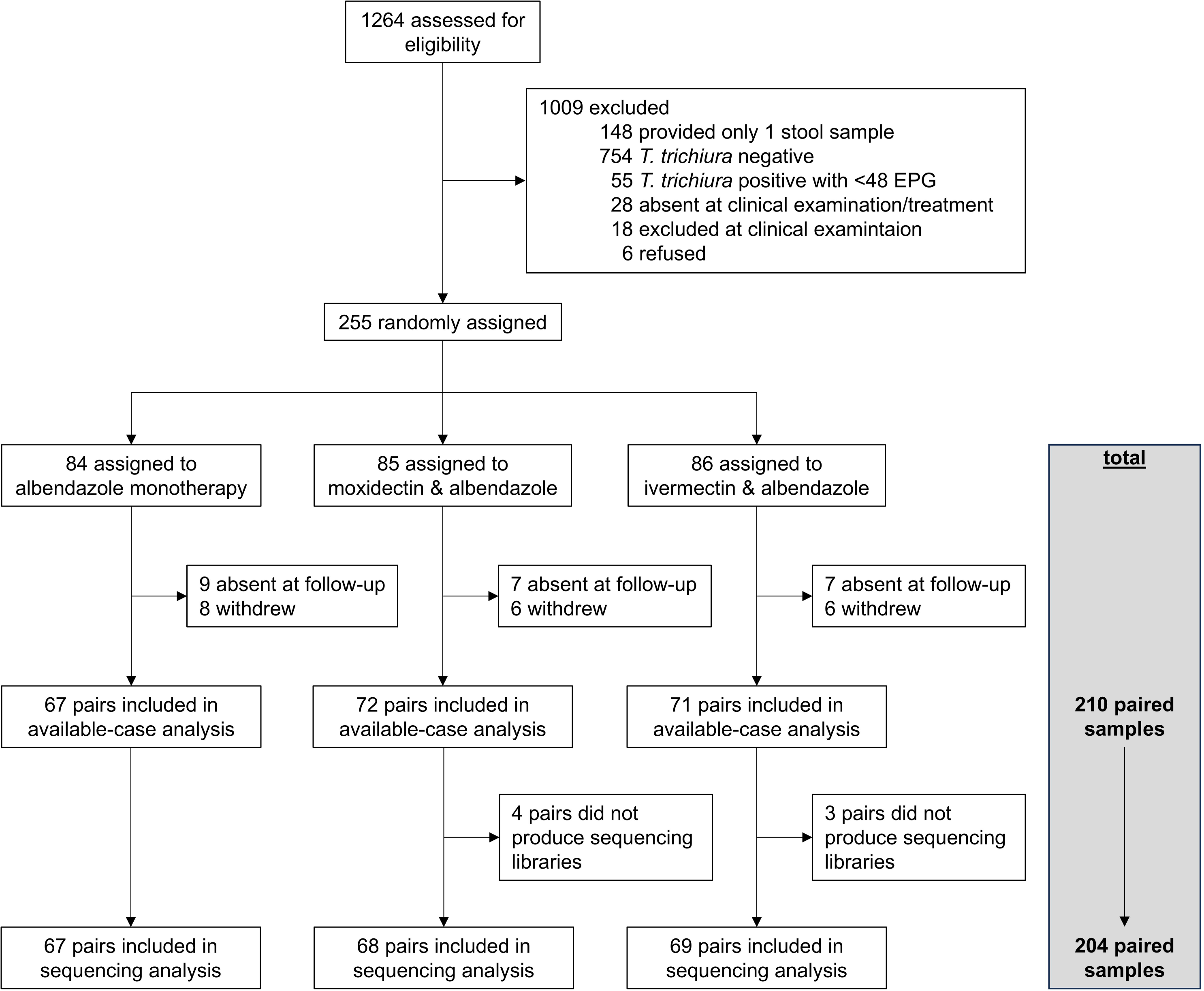
Trial profile (adapted from Sprecher et. al.^27^). Abbreviation: EPG, eggs per gram of stool.

**Table 1:**
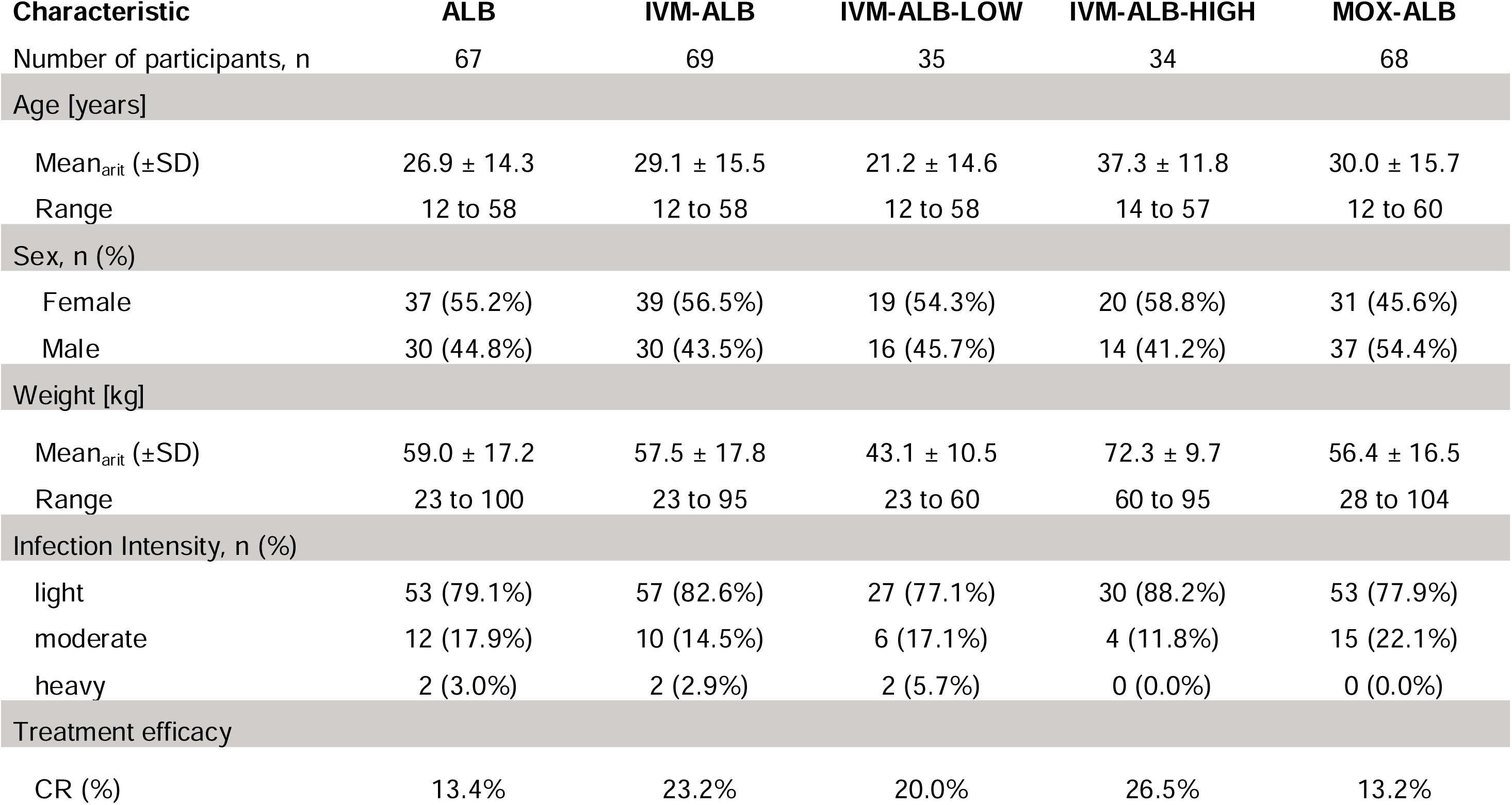
Study results summary, adapted from Sprecher *et. al.*^27^. Infection intensity classification is based on World Health Organization criteria^28^. Abbreviations: SD, standard deviation; CR, cure rate.

### DNA extraction of frozen stool samples

To extract DNA from frozen stool samples, we used the DNeasy PowerSoil Pro kit (QIAGEN, 47016) and followed the manufacturer’s standard protocol using 100-150mg of frozen stool. Final DNA concentrations were determined using a QuBit 4 fluorometer (Thermo Fisher Scientific, Q33238) with the 1x dsDNA HS Assay kit (Thermo Fisher Scientific, Q33231). Eluates were stored at −20°C until further use.

### Illumina and Nanopore sequencing

Illumina library preparation and sequencing were performed at the Genomics Facility of the Department of Biosystems Science and Engineering (D-BSSE) Switzerland. DNA eluates were adjusted to a concentration of 15-30ng/µl in nuclease-free water (Ultrapure™ Distilled water, Invitrogen, USA) prior to library preparation. Libraries were prepared using the NEBNext Ultra FS II (E7805L) reagents. Sequencing was performed on an S4 flow cell using an Illumina NovaSeq 6000 instrument targeting 18M paired-end reads per sample (2×150bp). Six samples did not produce any libraries, therefore six samples pairs were excluded from any further analyses (Figure 1). Nanopore libraries were prepared using the Oxford Nanopore Technologies Native Barcoding Kit 96 (SQK-NBD114-96) according to the manufacturer’s instructions. The libraries were sequenced on an Oxford Nanopore Technologies PromethION 2 solo instrument equipped with PromethION flow cells (FLO-PRO114M, R10.4.1) for 72h with a target of 3-5Gb sequenced bases per sample, according to Bertrand *et.al.* (avg. Q-score > 14, avg. reads over Q20 > 79%).

### Hybrid assemblies and binning

A comprehensive overview of this workflow is given in Supplementary Figure 1. We utilized Trimmomatic^29^ (v. 0.39) for adapter removal and quality filtering of the raw Illumina reads. The following setting were applied: -phred 33 ILLUMINACLIP:TruSeq3-PE-2.fa:2:30:10 LEADING:3 TRAILING:3 SLIDINGWINDOW:4:15 MINLEN:36. Subsequently, we removed non-bacterial reads utilizing Kraken 2^30^ (v. 2.1.3) equipped with the “MinusB” Refseq index (as of Oct 9^th^, 2023) and keeping the unclassified reads. Basecalling of the POD5 files obtained during Nanopore sequencing was performed using Dorado^31^ (v. 0.7.2) equipped with the model dna_r10.4.1_e8.2_400bps_sup@v4.3.0. We utilized Filtlong^32^ (v. 0.2.1) to filter short reads (< 300bp) and supplied the processed paired-end Illumina reads as a filtering template using the options “-1” and “-2” to retain only bacterial Nanopore reads. Lastly, adapters were removed using Dorado’s function “-trim”. Our aim was to generate eight pooled metagenomic assemblies representing the gut-bacterial microbiome of four treatment arms—ALB, IVM-ALB-LOW, IVM-ALB-HIGH, and MOX-ALB—at two timepoints: baseline (BL) and follow-up (FU). To begin, we prepared the Nanopore read data by generating subsets for each sample, capping them at 1 Gb using seqkit^33^ (v. 2.6.1) with the “seqkit head” command. This was necessary because the Nanopore sequencing depth varied between samples, ranging from 0.7 Gb to 14 Gb. By capping the data at 1 Gb, we ensured equal representation of long-read data from each sample in the pooled assemblies. Next, we pooled these Nanopore read subsets by treatment arm and timepoint. Illumina reads were also pooled by treatment arm and timepoint, but since their sequencing depth was consistent across samples, no subsetting was required. To finalize the datasets for assembly, we split each pooled dataset - both Nanopore and Illumina reads - into three separate subsets using seqkit commands (“seqkit split –p 3” for Nanopore and “seqkit split2 –p 3” for Illumina). This created technically independent triplicates for each of the eight pooled assemblies. These triplicates served two main purposes: to reduce computational resource demands and to identify and mitigate any technical artifacts that might arise during the assembly and binning processes. We started the assembly process by generating short-read assemblies via SPAdes^34^ (v. 4.0.0) via the command “spades.py –meta”, generating 24 metagenomic assemblies. We then utilized OPERA-MS^35^ (v. 0.9.0) equipped with the GTDB database^36^ (release 89 from June 17^th^ 2019) and using the long-read pools, as well as SPAdes short-read assemblies as input (via the flag “--contig-file”). Using the contigs put forth by OPERA-MS, we followed an ensemble binning approach utilizing MetaBAT2^37^ (v. 2:2.15), MaxBin2^38^ (v. 2.2.7) and CONCOCT^39^ (v. 1.1.0), followed by bin refinement with DAS Tool^40^ (v. 1.1.7). These bins underwent assessment in CheckM2^41^ (v. 1.0.2) and only bins with completeness > 95% and contamination < 5% were retained for further analyses. Clean, high-quality bins were taxonomically annotated with MetaPhlAn4^42^. A custom python script fragmented each bin into 150bp fragments that were then subjected to taxonomic annotation. Bins with ambiguous or no annotation were dropped from the analysis. To verify our annotation approach, we additionally annotated the bins using BAT^43^ (v. 6.0.1; Supplementary Table “SUPP_BAT_vs_MP4”). Since taxonomic annotation via BAT is stringent, we can only report matching taxonomies between the two techniques on a high taxonomic level. Subsequently, we utilized the species-level MetaPhlAn annotation and the CheckM2 reports to merge the bins put forth by the triplicate assemblies with a custom python script. Essentially, we kept all unique species-level bins, and only the most complete replicated bins (highest CheckM2 completeness) for each pool of bin triplicates in an arm and timepoint (Supplementary Figure 14).

### Phylogenetic trees

We constructed a phylogenetic tree representing both the BL and FU MAGs for each treatment arm using CheckM^44^ and the command “checkm tree”. As a quality control measure, we added a selection of identified core genomes of this study that were retrieved from the MetaPhlAn database, CHOCOPhlAn database (v. Oct22). The resulting alignment file was used as input for iqtree2^45^, for treebuilding and bootstrapping with the command “iqtree2 -m MFP -bb 1000 -alrt 1000 - nt 80”.

### Taxonomic profiling

We used the MetaPhlAn (v. 4.0.6) pipeline equipped with the CHOCOPhlAn database (v. Oct22) to compute taxonomic relative abundances for each sample, using paired-end Illumina reads as inputs. For subsequent analyses, we applied a 1% prevalence filter and a 20% variance filter in R-Studio (R-base v. 4.4.1) equipped with dplyr^46^ (v. 1.1.4). We then calculated alpha diversity measures utilizing the filtered relative abundance table. Richness, Shannon diversity indices, Berger-Parker dominance indices and Spearman correlations thereof were determined in R-Studio equipped with the following packages: car^47^ (v. 3.1-2), plyr^48^ (v. 1.8.9) and vegan^49^ (v. 2.6-6.1). To generate phylum-level relative abundance barplots, we used tidyr^50^ (v. 1.3.1) in addition. Bray-Curtis dissimilarities and non-Metric Multidimensional Scaling (NMDS) coupled to PERMANOVA and PERMDISP analyses were also facilitated by the package vegan (v. 2.6-6.1). The core microbiome was established via the microbiome^51^ (v. 1.28.0), phyloseq^52^ (v. 1.50.0) and reshape2^53^ (v. 1.4.4) packages. For differential abundance analysis of baseline and follow-up feature abundances and prevalences, we utilized MaAsLin3^54^ complemented with random forest analysis. For DAA, R-Studio was therefore additionally equipped with the following packages: maaslin3 (v. 0.99.1), randomForest^55^ (v. 4.7-1.2). In MaAsLin3 we the following model was used for each treatment arm: feature ∼ timepoint + (1 | subject_ID). Noteworthy, the unfiltered relative abundance table was used as input here, following the authors’ recommendations. Similarly, we applied the recommended settings for normalization (“TSS”) and transformation (“LOG”). For the illustrations, the R-packages RColorBrewer (v. 1.1-3), ggplot2^56^ (v. 3.5.1), ggsignif^57^ (v. 0.6.4), gridExtra (v. 2.3), ggpubr^58^ (v. 0.6.0), ggtext^59^ (v. 0.1.2), ggVennDiagram^60^ (v. 1.5.2) and cowplot^60^ (v. 1.1.3) were used. To generate the phylogenetic trees of the high-quality metagenomic assembled genomes (MAGs) for each arm, we utilized PhyloPhlAn3 (v. 3.1.68). To plot the obtained tree-files in R-Studio we made use of the following packages: ape^61^ (v. 5.8-1), ggtree^62^ (v. 3.14.0) and ggtreeExtra (v. 1.16.0).

### Analysis of functional profiles

The HUMAnN4^63^ pipeline (v. 4.0.0.alpha.1) equipped with the CHOCOPhlAn database (v. Oct22) was used to generate functional profiles of all samples based on the clean paired-end Illumina reads. Subsequently, we generated an unstratified table (merged taxonomy to pathway level) via the command “human_split_stratified_table” and converted the absolute numbers to relative abundances via “human_renorm_table -u relab”. This step was performed for both MetaCyc pathways (given by the “pathabundance” file) and MetaCyc reactions (given by the “genefamilies” file). Both relative abundance tables were subjected to 10% prevalence filtering in R. Analogous to the taxonomic relative abundance table, we calculated Bray-Curtis dissimilarities and performed NMDS coupled to PERMANOVA and PERMDISP analyses. Lastly, we also ran a differential abundance analysis on both relative abundance tables with MaAsLin3 and the model “feature ∼ timepoint + (1 | subject_ID)”. Both results were additionally filtered, keeping only the associations with the 20% lowest and 20% highest coefficients (for reactions) or the 5% lowest and 5% highest coefficients (for pathways; Supplementary Figures 22, 24). To compare the functional content of the pooled metagenomes of each arm, we annotated the MAGs using Bakta^64^ (v. 1.9.3). EC annotations of the MAGs were merged with a custom python script and plotted in R-Studio using packages described in this section previously.

### Analysis of resistome

To characterize the resistome in each treatment arm on reads-level (Figure 4) we used bowtie2^65^ (v. 2.5.2) equipped with a reference index comprised of the Comprehensive Antibiotic Resistance Database^66^ (CARD, v. 3.3.0), supplying the clean Illumina read files as mapping input. Subsequently, samtools (v. 1.19.2) with the command “idxstats” was used to compute the absolute number of mapped reads for each sample and reference ARG. We then translated the absolute count into relative counts normalized by gene count per million gene reads (GCPM). The relative abundance table was subjected to 1% prevalence filtering in R. Analyses of ARG beta-diversity and differential abundance analysis were then performed as described previously. We further characterized the resistome of the study cohort on MAG-level (Figure 5). We first used the Resistance Gene Identifier (RGI) equipped with the CARD database to detect ARGs in the MAGs. For each MAG, we subsequently generated a bowtie2 reference index comprised of its ARGs. Similarly to the read-level analysis, we then aligned the clean Illumina reads for all samples to these MAG-specific indices. Samtools (v. 1.19.2) with the command “idxstats” was used to compute the absolute number of mapped reads for each index. We then translated the absolute count into relative counts normalized by gene count per million and gene length (GCPM). Since we compared the fold changes of MAG-specific ARGs between timepoints to the fold change of the corresponding taxa, we annotated the MAGs with MetaPhlAn taxonomy. A custom python script fragmented each MAG into 150bp fragments that were then subjected to taxonomic annotation in MetaPhlAn. MAGs with ambiguous annotations or none were dropped from the analysis. Similarly, only taxa that were present both in BL and FU timepoints were taken forward for the analysis.

### Statistics and reproducibility

Group comparisons were conducted using Mann-Whitney U test for unpaired two-group comparisons and Wilcoxon Signed Rank-Sum tests for comparisons involving two paired groups, and all p-values were adjusted using the Benjamini-Hochberg correction – if not stated otherwise. To compare pairwise abundance and prevalence of taxa, MetaCyc pathways, MetaCyc reactions and ARGs we employed MaAsLin3 as described previously. For the subsequent analysis, we utilized the individual and joint “q-vals” put forth by MaAsLin3, which were also corrected using the Benjamini-Hochberg procedure. Both for MaAsLin3 and the PERMANOVA/PERMDISP analyses, we employed a significance threshold of p < 0.1. We provided a detailed documentation of the bioinformatics analysis (pipeline and R-scripts) under https://github.com/dommju/mac.

## Results

### Taxonomic profiling reveals treatment-specific shifts of gut-communities for the ALB, IVM-ALB-LOW, IVM-ALB-HIGH and MOX-ALB arms

We sequenced 67, 69 and 68 samples for the ALB, IVM-ALB and MOX-ALB arm, respectively (Figure 1). Paired-end Illumina sequencing yielded uniform read depths across all arms and timepoints, while Nanopore sequencing showed more variability (Figure 2A). For downstream analysis of the Nanopore data, 1Gb per sample was used (excluding three samples that did not meet this threshold). Diversity metrics (richness, Shannon’s H’, Berger-Parker Index (BP)) based on Illumina data showed no significant baseline differences across treatment arms (Supplementary Figure 2, Supplementary Table 1). Spearman correlations between timepoints indicated moderate consistency overall (*p* > 0.019), but richness and BP in the MOX-ALB arm did not correlate (richness: r_s_ = 0.44, *p* = 0.802; BP: r_s_ = 0.11, *p* = 0.379; Supplementary Figure 3). Phylum-level compositions remained stable over time (Wilcoxon tests: all *p* > 0.3; Supplementary Figure 4). However, species-level fold-change analysis revealed significant shifts in the MOX-ALB arm (*p* = 0.024), with lower follow-up prevalence of core species, except for *Anaerobutyricum soehngenii* (Supplementary Figure 5). No significant changes were observed in the ALB (*p* = 0.779) or IVM-ALB (*p* = 0.365) arms. Since IVM treatment is weight-based (200µg/kg), we further stratified the IVM-ALB arm based on the weight of the subject, utilizing a weight threshold of 60kg (IVM-ALB-LOW, n = 35; IVM-ALB-HIGH, n = 34). Diversity metrics remained unchanged between these subgroups at baseline (Supplementary Figures 6, Supplementary Table 1). Timepoint correlations were generally moderate to strong (r_s_ = 0.22-0.62, *p* < 0.05), except for H’ in the IVM-ALB-LOW group (*p* = 0.240; Supplementary Figure 7). Bray-Curtis dissimilarity showed modest but significant shifts between timepoints in ALB (*R²* = 0.0079, *p* = 0.032) and IVM-ALB arms (*R²* = 0.0059, *p* = 0.066), but not in the MOX-ALB arm or IVM-ALB subgroups (Figure 2B). PERMDISP analysis indicated significant dispersion only in the IVM-ALB-HIGH group (*F* = 2.896, *p* = 0.093), suggesting that the differences in the ALB and IVM-ALB arms are due to community-wide shifts and not dispersion thereof and just individual subjects in the IVM-ALB-HIGH arm experienced strong taxonomic shifts between timepoints.

**Figure 2:**
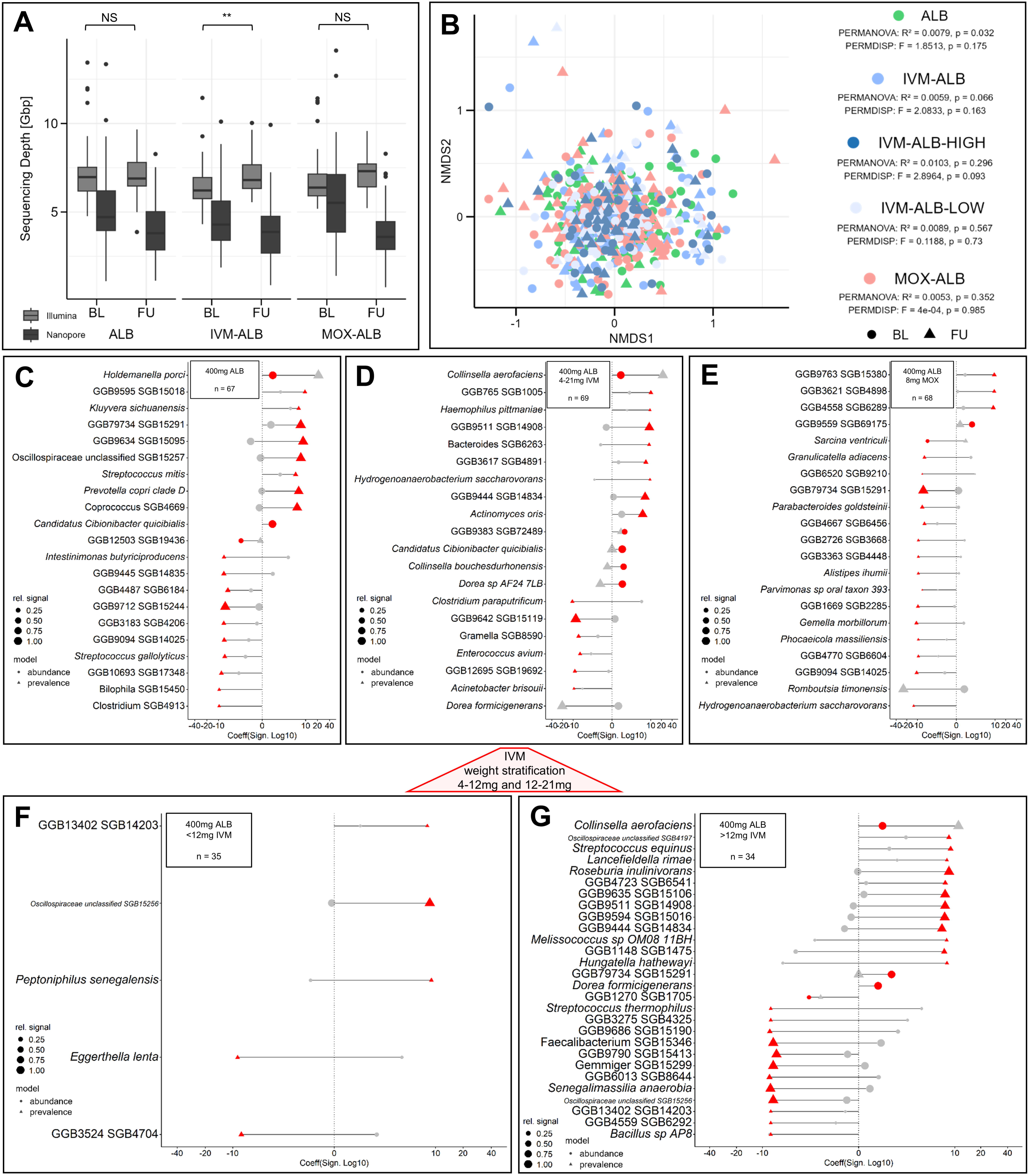
Taxonomic profiles for ALB, IVM-ALB and MOX-ALB cohorts before and after treatment. **A.** Sequencing depths for 408 samples both sequenced on the lllumina and Nanopore platforms. Boxplots show median (center line), interquartile range (box), 1.5xlQR whiskers, and outliers (dots). T-test (Bonferroni-corrected): *p* = 1 (ALB), 0.005 (IVM-ALB), 0.192 (MOX-ALB). **B.** Non-metric multidimensional scaling (NMDS) ordination based on Bray-Curtis Dissimilarity matrices of relative taxa abundances of paired BL and FU samples. The stratification of the IVM-ALB arm into the IVM-ALB-LOW and IVM-ALB-HIGH arms is based on a weight threshold of 60kg, corresponding to 12mg of IVM. PERMANOVA: *p* = 0.032 (ALB), 0.066 (IVM-ALB), 0.296 (IVM-ALB-HIGH), 0.567 (IVM-ALB-LOW), 0.352 (MOX-ALB). PERMDISP: *p* = 0.175 (ALB), 0.163 (IVM-ALB), 0.093 (IVM-ALB-HIGH), 0.730 (IVM-ALB-LOW), 0.985 (MOX-ALB). **C-G.** Pairwise, species-level abundance and prevalence modelling of taxa between timepoints reaching a "joint q-val" of < 0.1. Datapoints with "individual q-val" < 0.1 are depicted in red. Coefficients for the abundance models are given as logiFU/BL). Coefficients for the prevalence models are given as ln(OR). The relative signal was calculated as the ratio of samples, in which the association was detected divided by the total number of samples for that arm and timepoint. **C.** ALB **D.** IVM-ALB **E.** MOX-ALB **F.** IVM-ALB-LOW (weight< 60kg, IVM administered < 12mg). **G.** IVM-ALB-HIGH (weight> 60kg, IVM administered > 12mg).

Pairwise species-level modelling between timepoints (Figures 2C-G) revealed the most pronounced microbiome shifts in the IVM-ALB-HIGH arm (Figure 2G), with post-treatment enrichment of *Collinsella aerofaciens* (logl(FU/BL) = 0.8, *q* = 0.05), *Roseburia inulinivorans* (ln(OR) = 8.5, *q* = 0.03), and *Dorea formicigenerans* (logl(FU/BL) = 0.6, *q* = 0.08). Depletions were observed for *Faecalibacterium*-related species SGB15413 (ln(OR) = −6.8, *q* = 0.07), SGB15346 (ln(OR) = −7.4, *q* < 0.001), and *Senegalimassilia anaerobia* (ln(OR) = −8.0, *q* < 0.001). In contrast, the IVM-ALB-LOW arm (Figure 2E) showed enrichment of only one taxon, Oscillospiraceae-related SGB15256 (ln(OR) = 8.6, *q* < 0.001). The MOX-ALB arm (Figure 2F) was associated with enrichment of SGB69175 (logl(FU/BL) = 1.6, *q* = 0.08) and depletion of SGB15291 (ln(OR) = −6.7, *q* < 0.001). ALB treatment (Figure 2C) enriched *Holdemanella porci* (logl(FU/BL) = 0.8, *q* = 0.03), *Streptococcus mitis* (ln(OR) = 5.4, *q* = 0.06), and *Prevotella copri* (ln(OR) = 6.5, *q* < 0.001). Random forest analysis corroborated these trends, identifying *H. porci*, *C. aerofaciens*, *D. formicigenerans*, and SGB69175 as key discriminants between BL and FU communities (Supplementary Figures 8–12). Collectively, these results demonstrate that ALB and IVM-ALB-HIGH treatments elicit the strongest and most taxonomically diverse microbiome shifts, underscoring their specific yet broad impacts at the species level.

### Functional profiling of treatment-stratified, pooled metagenomes reveals profound differences between timepoints for the ALB and IVM-ALB-HIGH arms

We next assessed functional profiles across timepoints. Bray-Curtis dissimilarity of pathway abundances (Supplementary Figure 13) revealed small but significant differences between BL and FU for the ALB (*R²* = 0.0117, *p* = 0.035), IVM-ALB-LOW (*R²* = 0.0166, *p* = 0.098), and IVM-ALB-HIGH arms (*R²* = 0.0232, *p* = 0.005), with no change in the MOX-ALB arm (*R²* = 0.0049, *p* = 0.573). The strongest shifts occurred in IVM-ALB-HIGH, as reflected by the highest *R²* value. PERMDISP analysis confirmed significant dispersion in IVM-ALB-HIGH (*F* = 4.975, *p* = 0.027), indicating subject-level variation, while changes in the ALB and IVM-ALB-LOW arms reflected consistent group-wide shifts. Reaction-level beta-diversity (Figure 3A) followed a similar pattern, with significant differences in the ALB (*R²* = 0.0139, *p* = 0.024) and IVM-ALB-HIGH arms (*R²* = 0.0238, *p* = 0.015), but not in the IVM-ALB-LOW (*R²* = 0.0123, *p* = 0.332) or MOX-ALB arms (*R²* = 0.0049, *p* = 0.573). PERMDISP again suggested group-wide shifts in the ALB arm (*F* = 1.604, *p* = 0.176) versus subject-specific changes in the IVM-ALB-HIGH arm (*F* = 3.917, *p* = 0.053). These results indicate that IVM-ALB-HIGH treatment induces the most pronounced and individualized shifts in functional potential, both at the pathway and reaction levels.

**Figure 3:**
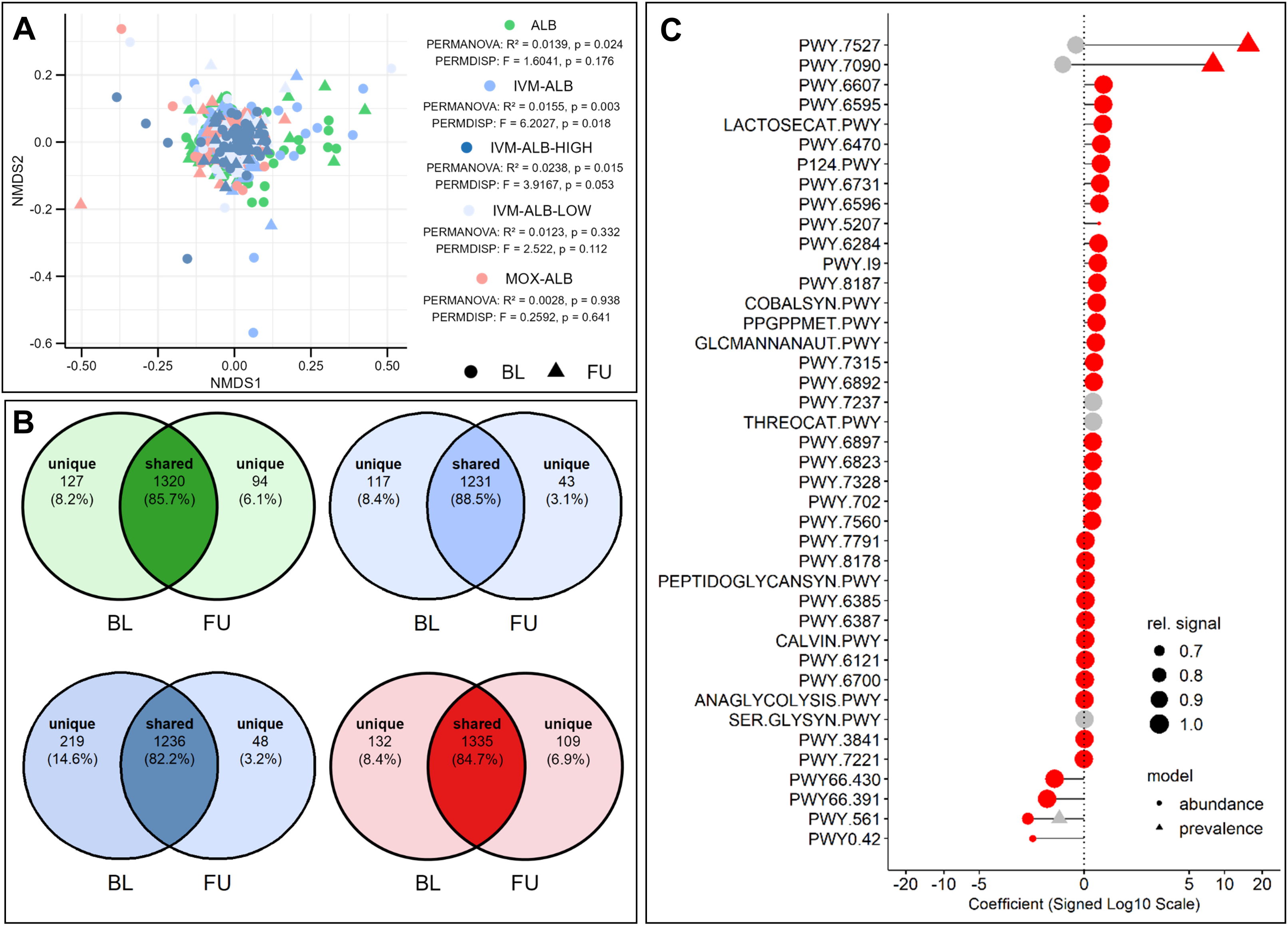
Functional profiles for ALB, IVM-ALB-LOW, IVM-ALB-HIGH and MOX-ALB cohorts before and after treatment. **A.** Non-metric multidimensional scaling (NMDS) ordination based on Bray-Curtis Dissimilarity matrices of relative MetaCyc reaction abundances of paired BL and FU samples. The stratification of the IVM-ALB arm into the IVM-ALB-LOW and IVM-ALB-HIGH arms is based on a weight threshold of 60kg, corresponding to 12mg of IVM. PERMANOVA: *p* = 0.024 (ALB), 0.003 (IVM-ALB), 0.015 (IVM-ALB-HIGH), 0.332 (IVM-ALB-LOW), 0.938 (MOX-ALB). PERMDISP: *p* = 0.176 (ALB), 0.018 (IVM-ALB), 0.053 (IVM-ALB-HIGH), 0.112 (IVM-ALB-LOW), 0.641 (MOX-ALB). **B.** Absolute and relative number of shared and unique enzyme commission (EC) numbers in metagenomic assembled genomes (MAGs) between BL and FU timepoints. **Top:** ALB (left), IVM-ALB-LOW (right). **Bottom:** IVM-ALB-HIGH (left), MOX-ALB (right). **C.** Pairwise, species-level abundance and prevalence modelling of pathways between timepoints of the IVM-ALB-HIGH arm after filtering, (keeping only the associations with the 20% lowest and 20% highest MaAslin3 coefficients and a relative signal > 0.5). The relative signal was calculated as the ratio of samples, in which the association was detected divided by the total number of samples for that arm and timepoint. Only pathways reaching a "joint q-val" of < 0.1 are shown. Pathways with "individual q-val" < 0.1 are depicted in red. Coefficients for the abundance models are given as log2(FU/BL). Coefficients for the prevalence models are given as ln(OR).

To contextualize functional shifts at the genome level, we analyzed metagenome-assembled genomes (MAGs) across timepoints. At BL, binning yielded 135, 92, 100, and 130 high-quality MAGs (completeness >95%, contamination <5%) for the ALB, IVM-ALB-LOW, IVM-ALB-HIGH, and MOX-ALB arms, respectively. At FU, we recovered 116, 95, 88, and 121 MAGs for the same arms. A substantial core of MAGs was shared across arms at each timepoint (43% at BL; 38% at FU), alongside smaller subsets unique to specific arms (7-16% at BL; 6-14% at FU; Supplementary Figure 15). Within-arm comparisons between timepoints revealed high persistence of MAGs in the ALB (49%), IVM-ALB-LOW (47%), and MOX-ALB arms (49%), indicating a stable core microbiome (Supplementary Figure 16), though each arm also showed ≥20% timepoint-specific MAGs. In contrast, the IVM-ALB-HIGH arm exhibited lower MAG overlap between timepoints (40%), with a higher proportion of unique MAGs at BL (34%) than FU (25%). To visualize these dynamics, we constructed treatment-specific phylogenetic trees integrating MAGs from both timepoints (Supplementary Figures 17-20), which showed broad phylogenetic diversity and several clades shared across timepoints in all treatment arms.

Gene content similarity between timepoints, assessed via EC annotation (Figure 3B), showed a consistently high overlap in enzymatic functions within each treatment arm (80-90%), indicating strong resilience of core microbiome functions to treatment. More pronounced functional differences between timepoints emerged from MaAsLin3-based pairwise comparisons of pathway abundance and prevalence (Figure 3C, Supplementary Figure 21). Filtering for pathways within the top and bottom 20% of model coefficients (Supplementary Figure 22) revealed four enriched at FU in the ALB arm: CDP-diacylglycerol biosynthesis I (PWY-5667), CDP-diacylglycerol biosynthesis II (PWY0-1319), starch degradation III (PWY-6731), and 1,5-anhydrofructose degradation (PWY-6992), all associated with phospholipid and carbohydrate metabolism. The IVM-ALB-HIGH arm exhibited the largest and most diverse set of enriched pathways at FU, spanning core metabolism (glycolysis III, pentose phosphate pathway, fatty acid β-oxidation VI), nucleotide turnover (guanosine and adenosine nucleotide degradation and biosynthesis), amino acid metabolism (e.g., L-arginine degradation XIII, L-cysteine biosynthesis VI), cofactor biosynthesis (e.g., molybdopterin, thiamine diphosphate), and stress responses (e.g., ppGpp metabolism). Enrichment of multiple peptidoglycan biosynthesis pathways (PWY.6387, PWY.6385, PEPTIDOGLYCANSYN.PWY) further indicated enhanced cell wall metabolism. Additional changes included the Bifidobacterium shunt and lactose/galactose degradation I. In the MOX-ALB arm, three FU-enriched pathways were observed, including two related to secondary metabolite biosynthesis (PWY-992, PWY-5910). No enriched pathways were detected in IVM-ALB-LOW. To further resolve these shifts, we examined reaction-level changes (Supplementary Figure 23), filtered to the top and bottom 5% of model coefficients (Supplementary Figure 24). In ALB, differentially abundant reactions were dominated by a single monooxygenase (EC 1.14.14.1), accounting for over 20 reactions. IVM-ALB-HIGH displayed broader enzyme class diversity, including transferases (EC 2.5.1.141, 2.7.1.207/208), dehydrogenases (EC 1.2.1.8, 1.1.1.130), hydrolases (EC 3.2.1.85, 3.1.13.5), and lyases (EC 4.4.1.16, 4.1.2.22). By contrast, IVM-ALB-LOW showed enrichment of a small set of oxidoreductases (EC 1.3.99.29, 1.17.7.1) and one hydrolase (EC 3.5.1.96), while MOX-ALB was associated with a lyase (EC 4.1.1.63) and two transferases (EC 2.1.1.230, 2.7.7.89). Taken together, these findings reveal nuanced, treatment-specific metabolic responses of the gut microbiome to these anthelmintic interventions, with IVM-ALB-HIGH inducing particularly strong and systemic functional reprogramming, likely driven by microbial restructuring or altered host–microbiota interactions.

### Resistome analysis of arm-specific metagenomic communities reveals mild differences between study timepoints

At BL, the gut resistome of the whole study cohort was dominated by ARGs targeting tetracyclines (56.1%, 84M reads), followed by those against streptogramins (7.1%, 11M), cephalosporins (6.2%, 9M), lincosamides (3.9%, 6M), and rifamycins (3.1%, 5M) (Figure 4A). Macrolide ARGs, relevant due to structural similarity to IVM and MOX, accounted for a smaller fraction (1.2%, 2M). ARG richness showed a modest increase at FU across arms, with only a few ARGs exceeding a richness threshold of >75 (Figure 4B). Across treatment arms, the majority of ARGs were shared between timepoints (ALB: 77.3%, IVM-ALB-LOW: 72.0%, IVM-ALB-HIGH: 73.4%, MOX-ALB: 83.0%), with smaller proportions unique to BL (7.5-15.8%) or FU (9.5-13.8%) (Figure 4C). ARG beta-diversity remained stable (Supplementary Figure 25), and most ARG classes were likewise conserved between timepoints (Figure 4D; shared: 81.7-86.7%, unique: BL 7.8-15.5%, FU 2.8-9.3%). Pairwise modelling of ARG dynamics (Figure 4E) revealed increased prevalence of the beta-lactamase *CfxA5* (ln[OR] = −9.3, *q* < 0.001) and decreased *tet(W/32/O)* (ln[OR] = −8.0, *q* < 0.001) in the IVM-ALB-LOW arm at FU. In the ALB arm, *CepA-44*, *CepA-49*, and *cepA* were more prevalent at BL (ln[OR] < −6.4, *q* < 0.001). Notably, the macrolide efflux pump *macB* increased significantly at FU in IVM-ALB-HIGH (ln[OR] = 8.0, *q* < 0.001), and was also more frequently detected in MOX-ALB at FU, though not significantly (ln[OR] = 11.8, *q* = 0.36).

**Figure 4:**
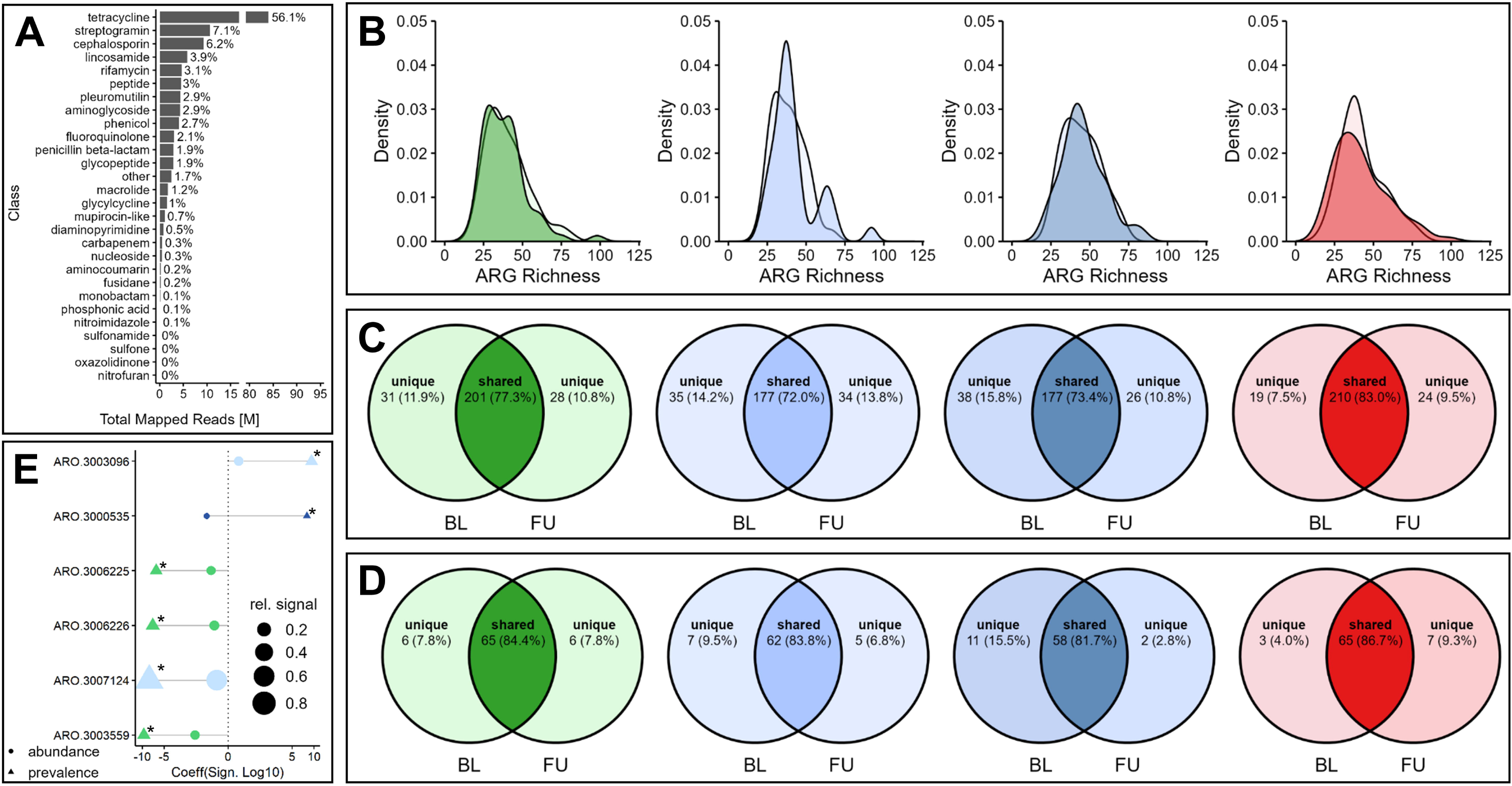
Resistome profiles based on lllumina sequencing reads for ALB, IVM-ALB-LOW, IVM-ALB-HIGH and MOX-ALB cohorts before and after treatment. **A.** Absolute abundance of antimicrobial resistance genes (ARGs) per antibiotic class in the whole study cohort at baseline (BL). **B.** ARG richness for BL and follow-up (FU) cohorts represented in a density function. **Left to right:** ALB, IVM-ALB-LOW, IVM-ALB-HIGH, MOX-ALB. Darker colors represent the FU content, lighter colors represent the BL content. **C-D.** Absolute and relative number of shared and unique ARGs **(C)** and ARG classes **(D)** between BL and FU timepoints. Left to right: ALB, IVM-ALB-LOW, IVM-ALB­ HIGH, MOX-ALB. **E.** Pairwise abundance and prevalence modelling of ARGs between timepoints reaching a "joint q-val" of< 0.1. Datapoints with "individual q-val" < 0.1 are marked with "*". Coefficients for the abundance models are given as log_2_(FU/BL). Coefficients for the prevalence models are given as ln(OR). The relative signal was calculated as the ratio of samples, in which the association was detected divided by the total number of samples for that arm and timepoint. Green= ALB, light-blue= IVM-ALB-LOW, dark-blue= IVM-ALB-HIGH.

To contextualize resistome shifts at a genome-resolved level, we examined ARG dynamics within high-quality MAGs (>95% completeness, <5% contamination) across timepoints, comparing mean ARG abundance to that of their corresponding host taxa (Figures 5A-D). Overall, ARG changes closely mirrored host MAG dynamics, suggesting limited gene-specific selection, with only a few exceptions (residual >2): In the ALB arm (Figure 5A), *vanT-G* (*Ruminococcus bromii*) and *vanY-B* (SGB4183) deviated from this trend. In the IVM-ALB-LOW arm (Figure 5B), several vancomycin resistance genes - *vanY-A* (*Turicibacter bilis*), *vanW-I*, and *vanT-G* (both from *Lachnospiraceae* spp.) - also diverged. *VanY-A* was disproportionately enriched at FU relative to its host MAG, whereas *vanW-I* and *vanT-G* were primarily associated with BL, despite balanced host abundance. A similar pattern was observed for *AAC(6’)-Iid* (*Enterococcus hirae*). In the IVM-ALB-HIGH arm (Figure 5C), *vanT-G* (SGB5099) showed increased abundance at FU, while its parent MAG was more frequent at BL, further indicating gene-level decoupling from taxon-level trends. Lastly, in the MOX-ALB arm, *vanT-G* and *vanY-B* exhibited disproportionately lower abundance relative to their parent MAGs, *Clostridium perfringens* and *Erysipelotrichaceae bacterium 7770 A6*, respectively. In the ALB arm, taxa with strong abundance shifts between timepoints (|loglFC| > 2) included *Akkermansia muciniphila* (carrying *adeF*), *Comamonas kerstersii* (*adeF*, *qacG*), and *Desulfovibrio piger* (*qacG*), all enriched at BL. While *Blautia stercoris* abundance was also higher at BL, its associated *poxtA* gene was more abundant at FU. In both IVM-ALB-LOW and IVM-ALB-HIGH arms, *Romboutsia timonensis* and *Turicibacter bilis* increased at FU, accompanied by elevated levels of *tet36* and vancomycin resistance genes (Figures 5F-G). Lastly, in the MOX-ALB arm, *Clostridium perfringens* and *Desulfovibrio piger* were enriched at BL, along with *mprF* and vancomycin resistance genes. At FU, vancomycin resistance genes varied in abundance and were coupled to increased levels of *Phocaeicola vulgatus*, *Roseburia faecis*, *Ruminococcus* spp., and *Streptococcus salivarius*. In summary, while specific ARGs - such as vancomycin resistance genes - show timepoint-specific deviations in select taxa, the cumulative gut resistome remains largely stable in response to treatment across all arms.

**Figure 5:**
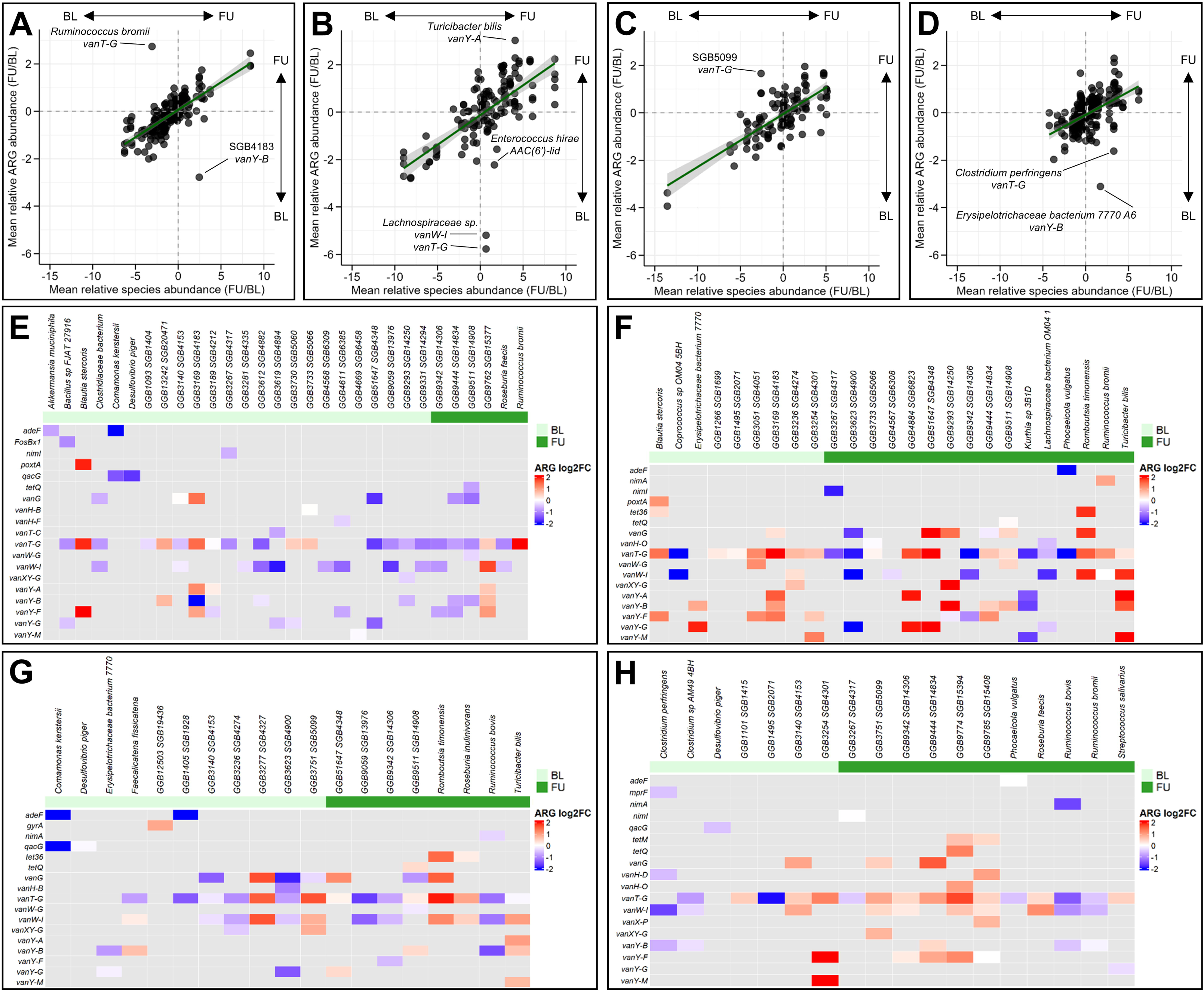
Resistome profiles of metagenomic assembled genomes for ALB, IVM-ALB-LOW, IVM-ALB-HIGH and MOX-ALB cohorts before and after treatment. **A-D.** Scatter of mean log_2_-transformed antimicrobial resistance gene (ARG) abundance fold-change originating from metagenomic assembled genomes (MAGs) and the corresponding mean log_2_-transformed species abundance fold-change from MetaPhlAn profiles (Figure 1) between the baseline (BL) and follow-up (FU) timepoints (FU/BL). The corresponding species abundances were matched with the ARG abundances by applying the MetaPhlAn taxonomy to the MAGs. **A.** ALB **B.** IVM-ALB-LOW **C.** IVM-ALB-HIGH **D.** MOX-ALB. **E-H.** Mean log_2_-transformed antimicrobial resistance gene (ARG) abundance fold-changes (FU/BL) for the MAGs exerting a mean log_2_-transformed species abundance fold-change (FU/BL) > 2 or < −2. We used the existing MetaPhlAn profiles (Figure 1) to calculate mean abundance fold-changes (FU/BL) for each species and subsequently applied the MetaPhlAn taxonomy to the MAGs. **E.** ALB **F.** IVM-ALB-LOW **G.** IVM-ALB-HIGH **H.** MOX-ALB.

## Discussion

Given the limited arsenal of anthelminthic treatment options, combination therapies involving IVM and MOX offer a well needed perspective for a better treatment option. Beyond their well-established roles in treating parasitic infections, IVM in particular is investigated for malaria transmission control through high-dose regimens^67^. However, despite this promising potential, current evidence raises several concerns. First, the efficacy of macrocyclic lactone combination treatments against STH infections exhibits notable geographical and species-specific variability^27,68^. Moreover, recent *in vitro* studies have highlighted the broad-spectrum antibacterial properties of these compounds^22,23^. Yet, comprehensive data on their potential off-target effects in human populations remain scarce.

For IVM, previous investigations on the gut microbiome have been constrained in terms of sample size, taxonomic resolution (genus-level), and the use of different partner drugs, such as tribendimidine, as well as by the lack of direct comparison to the current standard of care (e.g. ALB)^69^. In this study, we address these gaps by providing the first in-depth characterization of off-target taxonomic and functional changes following ALB, IVM-ALB or MOX-ALB treatment in a large cohort of *T. trichiura*-infected individuals in Côte d’Ivoire. We employ a state-of-the-art hybrid shotgun sequencing approach combined with advanced bioinformatics workflows to enable genome-resolved analyses of changes in the gut microbiome associated with exposure to different anthelmintics, at both the taxonomic and functional levels. We further extend the public health relevance of this work by examining the modulation of the gut resistome across more than 800 high-quality metagenomes.

From a taxonomic perspective, we did not observe high-level changes, such as shifts in the relative abundance of phyla or the prevalence of core microbial species. However, pairwise comparisons at the species-level revealed significant changes in abundance and prevalence between timepoints across all treatment arms. These findings align with existing literature, which describes the taxonomically broad yet species-specific antibacterial properties of both IVM and MOX that are also concentration-dependent^22,23^. A key pharmacological difference between the two anthelminthics lies in their administration: MOX is administered at a fixed dose of 8 mg, whereas IVM dosing is weight-based (200 µg/kg). According to Maier *et. al.* estimated intestinal IVM concentrations for subjects of this trial range from 1.8µM to 7.2µM and according to recent *in vitro* data IVM inhibits growth of several gut-bacterial isolates in concentrations ≥ 5µM^23^. Indeed, the stratification of the IVM-ALB cohort by weight - into IVM-ALB-LOW (<12 mg) and IVM-ALB-HIGH (>12 mg) arms - revealed a markedly greater extent of off-target effects in the IVM-ALB-HIGH arm. This stratification introduced the risk of age bias; however, the age ranges were similar between arms (IVM-ALB-LOW: 12–58 years; IVM-ALB-HIGH: 14–57 years), and BL composite microbiome metrics did not differ significantly. Therefore, increased total amount of drug administered, and consequently increased intestinal drug concentrations, likely account for the observed off-target effects in the IVM-ALB-HIGH arm. This interpretation is further supported by data from the IVM-ALB-LOW arm, where the taxonomic signals associated with treatment were comparably sparse. Currently, there are no data for estimated intestinal concentrations of MOX, but despite its lower molecular weight (IVM: 875.1g/mol, MOX: 639.8g/mol), we argue that the intestinal concentrations of MOX in the MOX-ALB cohort are lower compared to IVM in the IVM-ALB-HIGH cohort given the difference in administered dose: Every individual in the IVM-ALB-HIGH arm received a dose exceeding 8 mg, with 9 out of 34 participants receiving more than 16 mg of IVM. This fact likely explains the sparsity in detected off-target effects in the MOX-ALB arm, similar to the IVM-ALB-LOW arm and in stark contrast to the IVM-ALB-HIGH arm.

This trend becomes even more pronounced at the functional level and is consistently confirmed across multiple analytical approaches. First, ordination analyses revealed significant differences between timepoints in the IVM-ALB cohort, while no such effect was observed for the MOX-ALB cohort. Notably, within the IVM-ALB-HIGH arm, a substantially larger proportion of variance was explained by timepoint, regardless of the functional resolution—whether at the level of individual reactions or broader metabolic pathways. In the IVM-ALB-HIGH arm, the observed shifts in ordination centroids are likely driven by increased beta dispersion. This suggests that functional shifts are primarily occurring within individual participants, rather than reflecting a uniform, community-wide transformation. Supporting this interpretation, pairwise modelling of reaction and pathway abundance and prevalence revealed widespread modulation across numerous functional domains in response to IVM-ALB-HIGH treatment, whereas no comparable effects were detected in the IVM-ALB-LOW or MOX-ALB arms. The modulation of functional profiles in the IVM-ALB-HIGH arm indicates substantial remodeling of community structure (as observed by taxonomy) and metabolism, characterized by the excessive proliferation and elimination of specific microbial species, as well as alterations in the synthesis and degradation of cellular structures and metabolites, with particular focus on peptidoglycan biosynthesis. Interestingly, the trend of profound functional modulation did not extend to ARGs; across all cohorts, we did not detect significant associations between timepoints and the abundance of ARGs. However, a noteworthy exception was *macAB*, an efflux pump known to confer resistance to macrolide antibiotics among other classes. This resistance gene was found to be enriched at the FU timepoint in the IVM-ALB-HIGH arm, potentially indicating a concentration-dependent induction of macrolide resistance mechanisms at higher IVM concentrations – and – giving species with such pre-existing, broad resistance mechanisms a selective advantage when exposed to IVM.

In contrast to the macrolides IVM and MOX, current literature does not conclusively report antibacterial properties for ALB. Previous studies investigating off-target effects of single-dose ALB in a cohort of 32 hookworm-infected individuals from Ghana linked successful treatment to significant structural and compositional alterations in the gut microbiota^70^. Notably, such taxonomic changes were absent in individuals where treatment failed. These findings suggest that observed microbiota shifts are more likely attributable to microbial rearrangements following helminth clearance, rather than direct effects of ALB itself. This interpretation is further supported by existing evidence indicating strong associations between helminth infection status, anthelminthic treatment efficacy, and gut microbiota composition^71–73^. In our study, however, the low cure rates for *T. trichiura* infections suggest that the observed microbiota shifts in the ALB cohort are likely uncoupled from infection clearance or parasite species. Instead, these shifts may represent either indirect effects or previously underappreciated direct impacts of ALB treatment. While we observed multiple enriched and depleted taxonomic features in the ALB-treated cohort, the functional profiles revealed only minimal differences between timepoints, confined to a limited set of metabolic functions (e.g., EC 1.14.14.1). Furthermore, although ordination analyses detected significant taxonomic and functional shifts, the proportion of variance explained by timepoint was low. This indicates that the observed centroid shifts likely result from weak, diffuse community-wide changes rather than robust functional reprogramming. Importantly, as we did not consistently detect shared taxonomic or functional signatures attributable specifically to ALB across treatment arms, we propose that these signals represent unsystematic species turnover within the ALB cohort, with no measurable biological consequence. Nevertheless, we cannot entirely exclude the possibility that ALB induces some level of gut microbiota modulation. Moreover, it is plausible that the off-target effects measured in the IVM-ALB and MOX-ALB cohorts are at least partially influenced by the concurrent administration of ALB.

In summary, we demonstrate that MOX-ALB, as well as IVM-ALB at lower concentration, do not induce substantial off-target alterations in the gut microbiota. Similarly, ALB-associated microbiota shifts appear limited in scope and likely represent unsystematic species turnover rather than biologically meaningful community remodeling. In contrast, higher amounts of IVM (≥12 mg) were associated with pronounced taxonomic and functional perturbations, underscoring a concentration-dependent relationship in off-target effects, parallel with antibiotic exposure. These findings suggest that previously reported antibacterial properties of macrocyclic lactones may be more nuanced and context-dependent than generally assumed, influenced by dosage, treatment regimens, and host-specific factors. From a public health perspective, these results are especially relevant given the increasing interest in using higher-dose IVM regimens in malaria vector control programs^67^ and in treating STH infections^74^, especially when targeting adult populations. While such strategies may enhance antiparasitic efficacy or transmission interruption, our findings highlight the need to consider potential microbiota-mediated consequences, which could impact gut health, immunity, or pathogen resistance dynamics in treated communities.

Despite the added value of our findings, we acknowledge several limitations of this study. First, our results are derived from a limited geographic location—7 villages in Côte d’Ivoire—while *T. trichiura* infections are widespread in the southern hemisphere. Despite a potentially lower bias of parasite clearance in our study due to low cure rates, host-microbiome interactions vary across different populations and the generalizability of our findings to other geographical and demographical settings remains uncertain. Lastly, having increased longitudinal sampling points might help to further refine the off-target effects associated with anthelmintic exposure, especially long-term follow-ups at 6 to 12 months. This would help distinguish between transient and persistent effects.

### Ethical approval and consent to participate

The clinical study was conducted in compliance with the Declaration of Helsinki and International Council of Harmonization Good Clinical Practice guidelines and approved by the responsible ethics committees in Switzerland and Côte d’Ivoire, as well as the Ivorian drug regulatory authority. The clinical trial was registered under the number NCT04726969 on ClinicalTrials.gov. All study participants were informed about the purpose of the study and informed consent forms were obtained during information sessions.

## Supporting information

SUPP_BAT_vs_MP4

## Data Availability

The sequencing data (ONT and Illumina shotgun sequencing) generated in this study have been deposited in the NCBI Short Read Archive under the accession PRJNA1249224 (Reviewer link: https://dataview.ncbi.nlm.nih.gov/object/PRJNA1249224?reviewer=uhn20ea5ne6ei6iofvk9nuesqr). Numerical source data underlying all figures, as well as detailed documentation of the bioinformatics analysis can be found under https://github.com/dommju/mac.

## Data Availability for Clinical Trials

The clinical trial was registered under the number NCT04726969 on ClinicalTrials.gov. The results of the trial were published by Sprecher *et al*.^27^.

## Code Availability

The detailed documentation of the bioinformatics analysis can be found under https://github.com/dommju/mac.

## Author Contributions

J.D.: study design, research design, project supervision, experimental work, statistical analyses, figure generation, writing of the initial paper, and paper editing. V.P.S.: study design, research design, project supervision, field implementation, and paper editing. C.B.: experimental work (Illumina sequencing), paper editing. D.B.: experimental work (Nanopore sequencing), paper editing. E.H.: study design, research design, project supervision, field implementation, and paper editing. J.T.C.: study design, research design, project supervision, and paper editing. J.K.: study design, research design, project supervision, funding acquisition, and paper editing. P.H.H.S.: study design, research design, project supervision, funding acquisition, and paper editing.

## Acknowledgements

We would like to thank all participants of the trial (NCT04726969) for their trust and interest in participating. We further want to express our gratitude towards the health workers of the 7 villages in Dabou and Jacqueville sanitary districts, as well as all trial team members of the Centre Suisse de Recherches Scientifiques en Côte d’Ivoire, Université Félix Houphouët-Boigny in Abidjan and the Dabou Methodist Hospital, Côte d’Ivoire for their tireless effort. All steps of the computational analysis were performed using the scientific computing centre at the University of Basel (sciCORE; http://scicore.unibas.ch/). We are grateful to the European Research Council (No. 101019223) for financial support.

## Declaration on the use of AI

In the process of preparing this manuscript, AI-tools (ChatGPT) were leveraged for language editing, and grammatical correction. During preparation of the GitHub repository, AI-tools (ChatGPT) were utilized for code clean-up and code annotation (R-scripts). Hence, these tools solely served an editorial purpose and did not contribute to the study’s concept, analysis or conclusion. All scientific content in this manuscript represents the sole responsibility of the authors, who have thoroughly reviewed and approved the final versions of the text and illustrations.

