## Supplementary Material for "Hybrid-metagenomics reveal off-target effects of albendazole, ivermectin-albendazole and moxidectin-albendazole on the human gut bacteria"

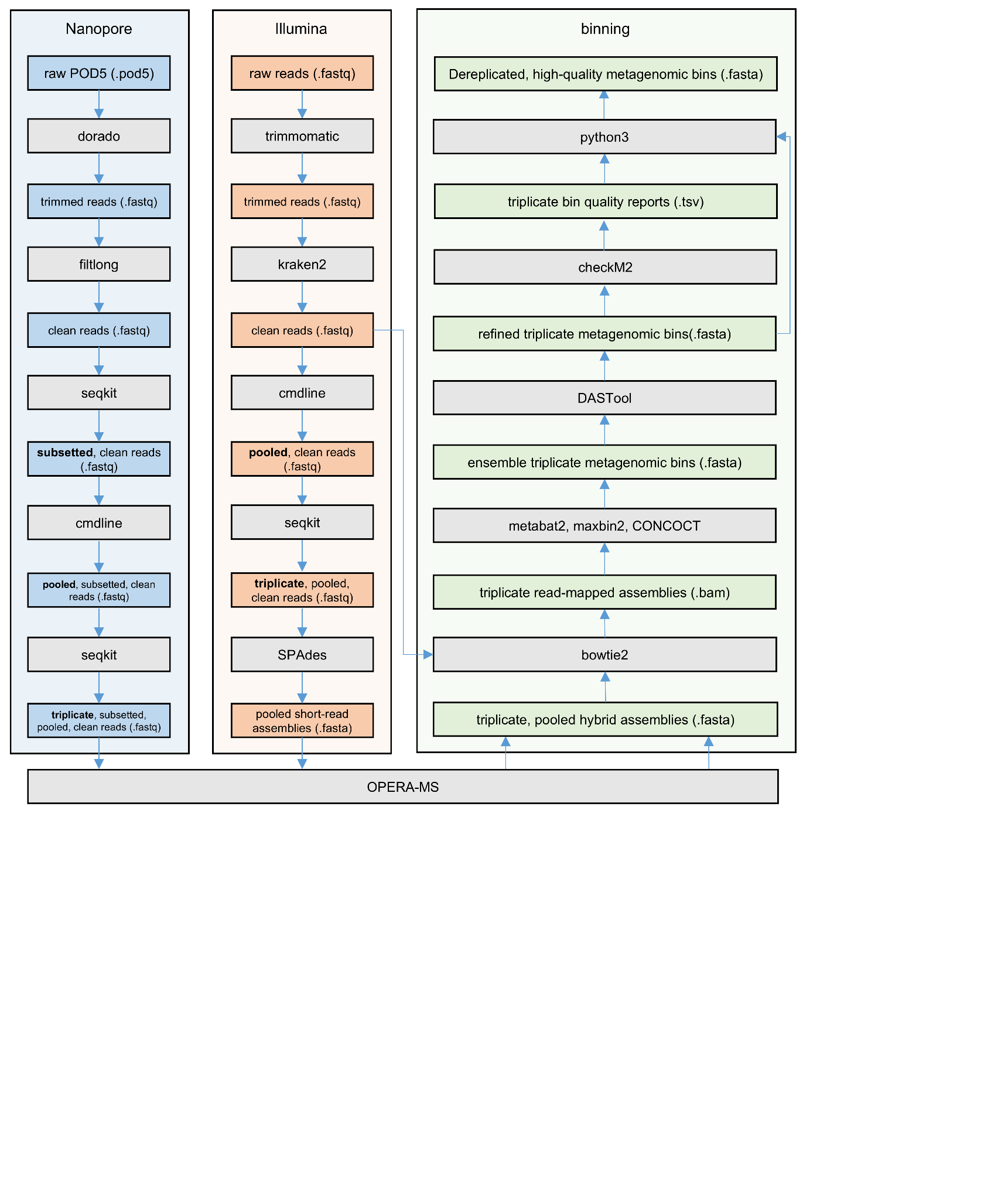

**Supplementary Figure 1:** Bioinformatics pipeline utilized to process the raw sequencing files from both the Illumina and Nanopore platform to form high-quality metagenomic bins.

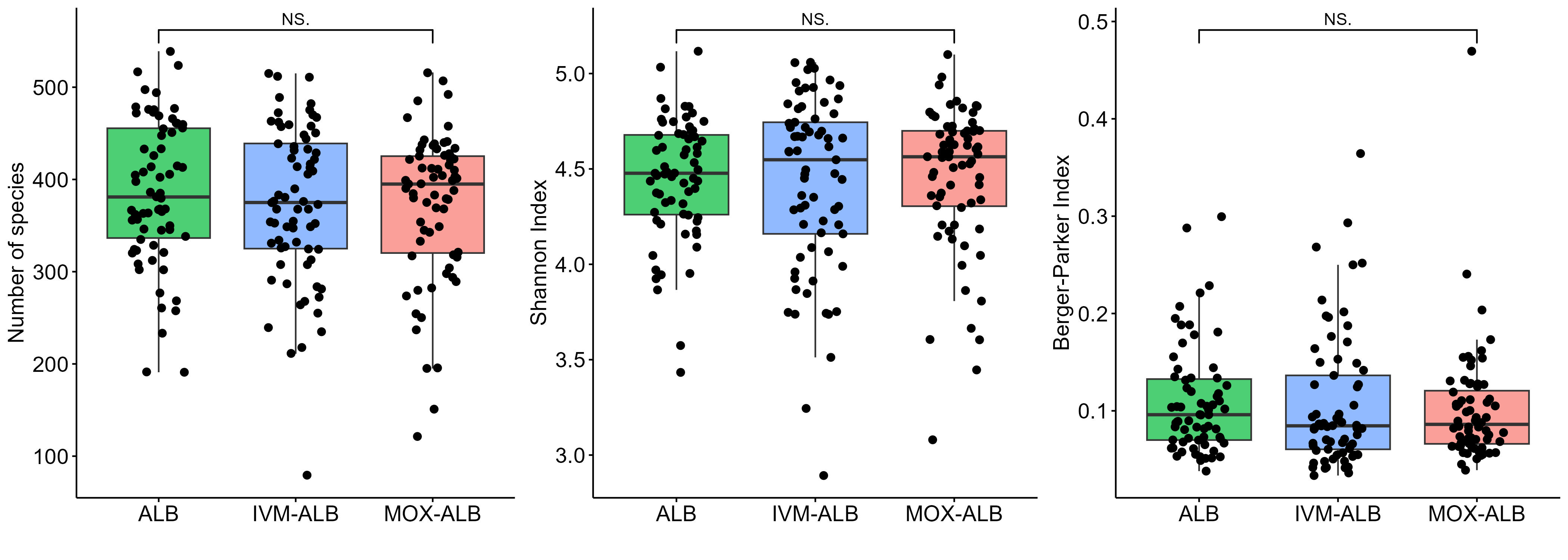

**Supplementary Figure 2:** Distributions of the composite metrics richness, Shannon Index and Berger-Parker Index in the baseline cohort, stratified by treatment arm. We utilized a Wilcoxon Rank Sum test to assess statistical significance.

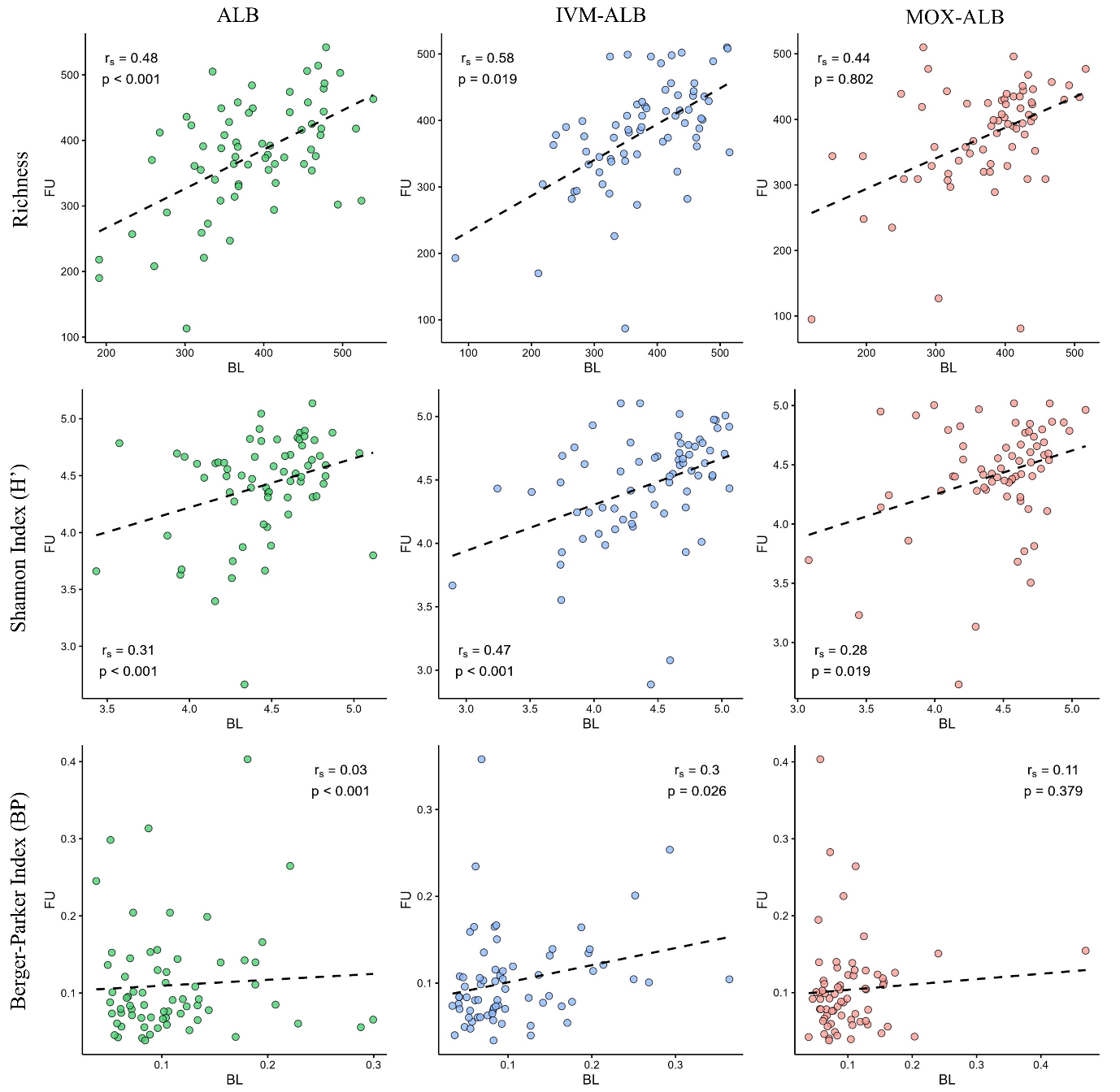

**Supplementary Figure 3:** Spearman correlations of the composite metrics Richness, Shannon Index and Berger-Parker Index for baseline (BL) and follow-up (FU) timepoints in each treatment arm.

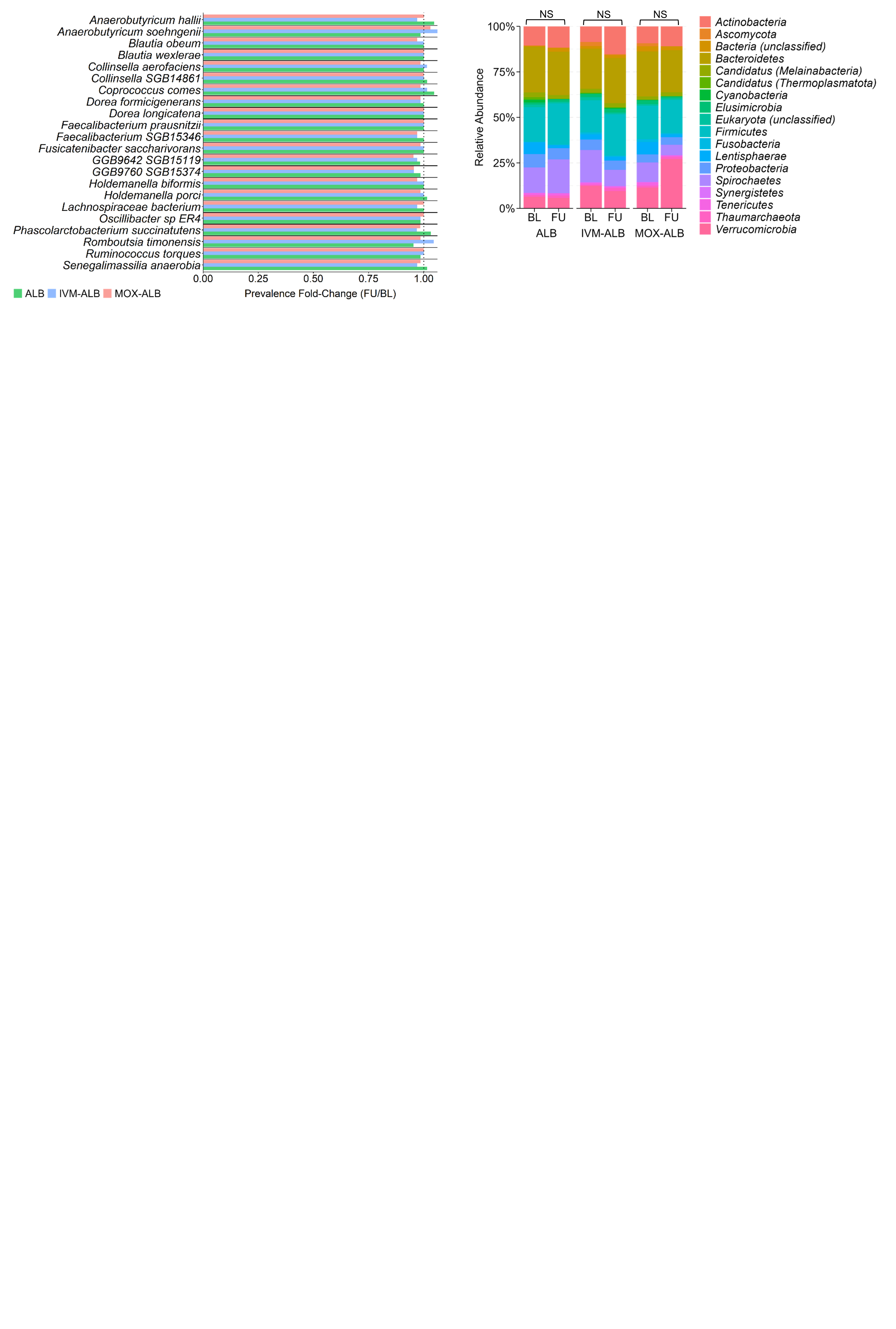

**Supplementary Figure 4:** Phylum-level relative abundances for baseline (BL) and follow-up (FU) timepoints across three treatment arms ALB, IVM-ALB and MOX-ALB. Wilcoxon Signed-Rank Test (BH-corrected): p = 0.832 (ALB), 0.832 (IVM-ALB), 0.389 (MOX-ALB).

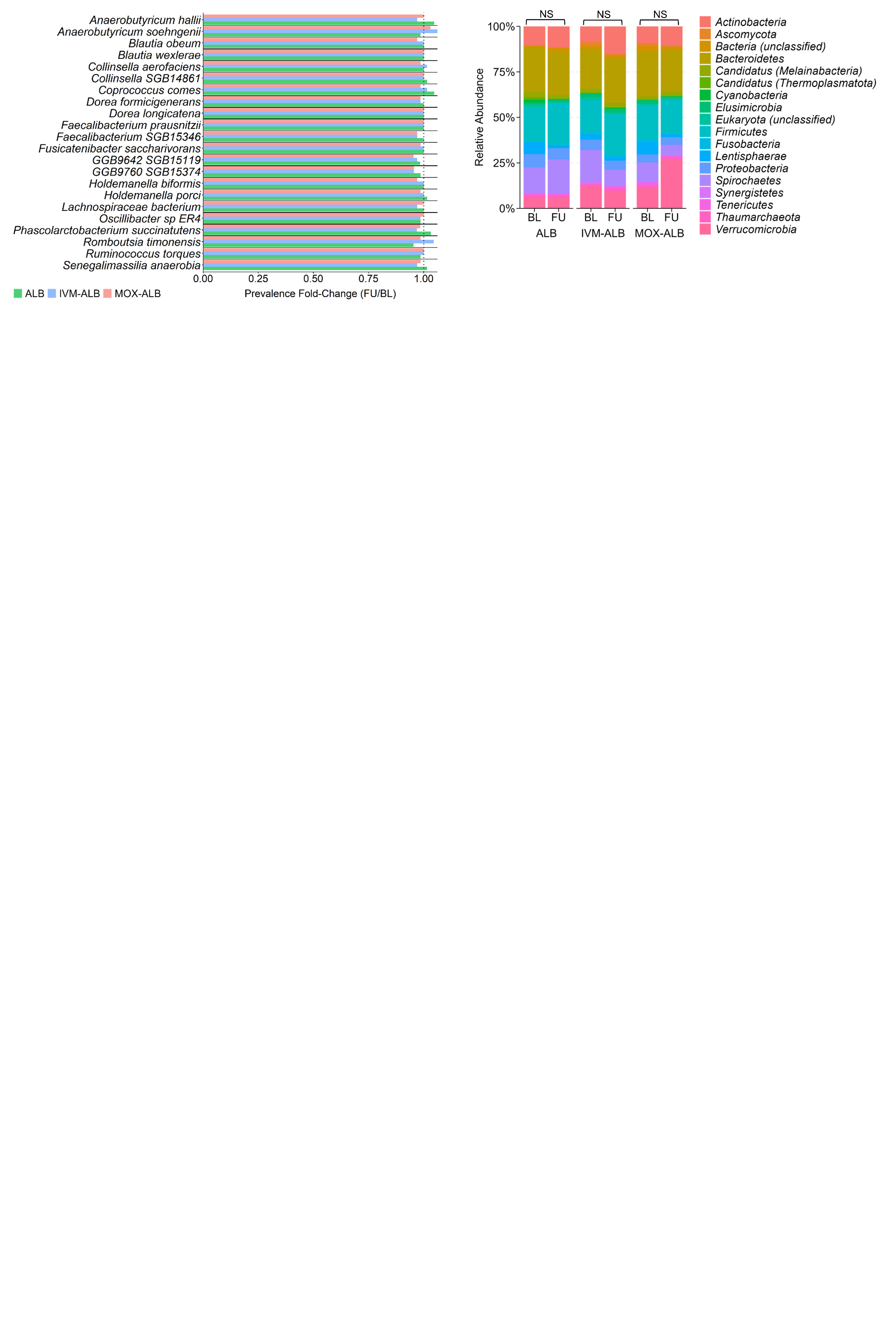

**Supplementary Figure 5:** Prevalence fold-change of core microbiota (FU/BL) for the three treatment arms ALB, IVM-ALB and MOX-ALB. Wilcoxon Rank-Sum Test (BH-corrected): p = 0.779 (ALB), 0.365 (IVM-ALB), 0.024 (MOX-ALB).

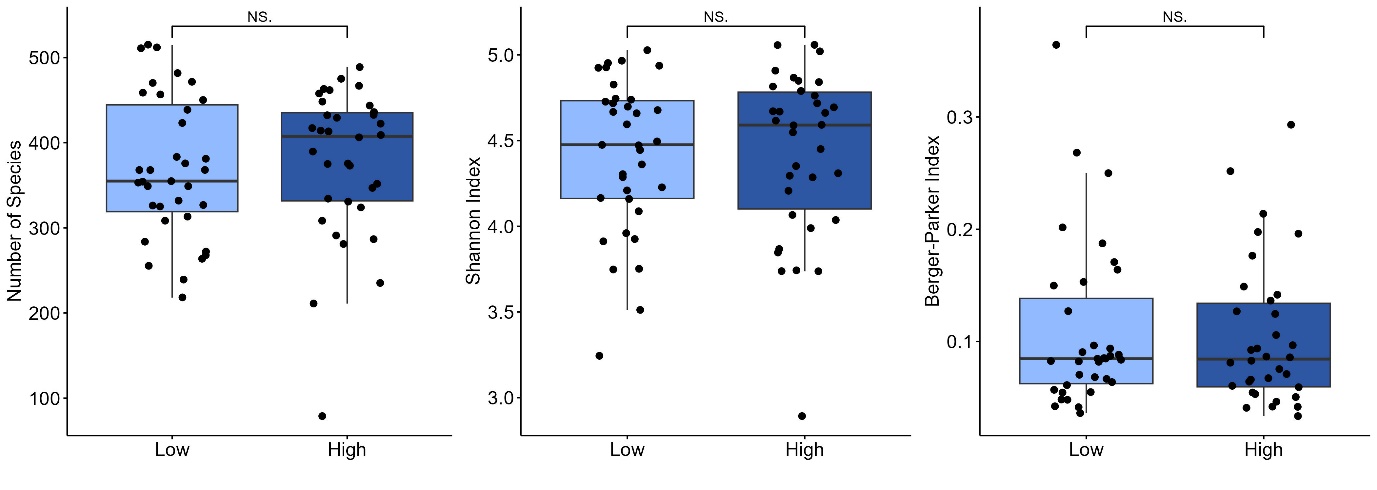

**Supplementary Figure 6:** Distributions of the composite metrics richness, Shannon Index and Berger-Parker Index in the baseline cohort of the IVM-ALB arm, stratified by dosage category. We utilized a Wilcoxon Rank Sum test to assess statistical significance.

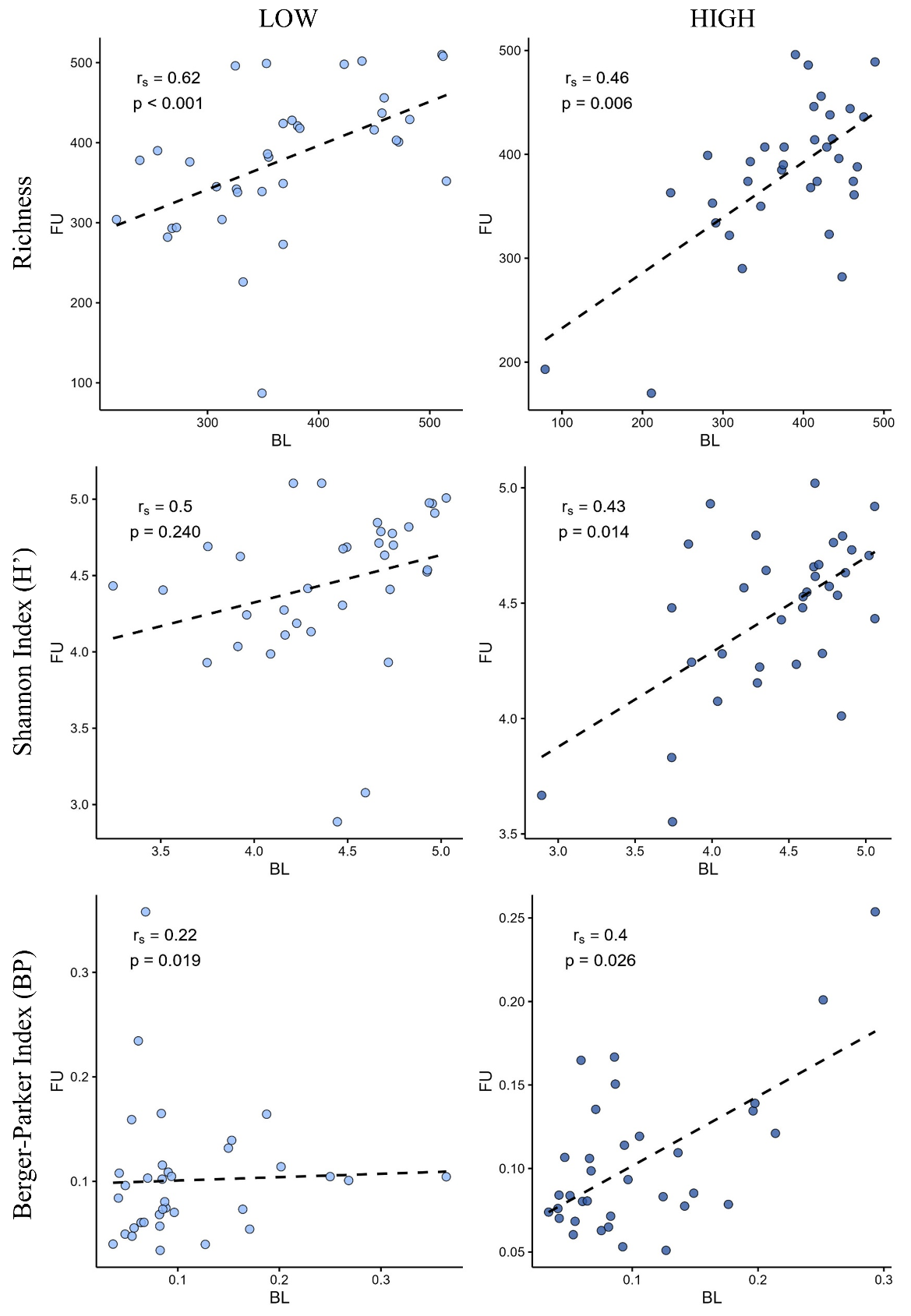

**Supplementary Figure 7:** Spearman correlations of the composite metrics richness, Shannon Index and Berger-Parker Index for baseline (BL) and follow-up (FU) timepoints in the two dosage categories of the IVM-ALB arm.

**Supplementary Table 1:** P-values of Mann-Whitney Test comparing baseline composite metrics (Shannon Diversity (H’), richness and Berger-Parker Index (BP)).

| **p-value (BH)** | **MOX-ALB vs. ALB** | **IVM-ALB vs. ALB** | **IVM-ALB-LOW vs. IVM-ALB-HIGH** |
| --- | --- | --- | --- |
| **H'** | 0.7659 | 0.8909 | 0.8909 |
| **richness** | 0.7659 | 0.7659 | 0.7659 |
| **BP** | 0.7659 | 0.7659 | 0.8909 |

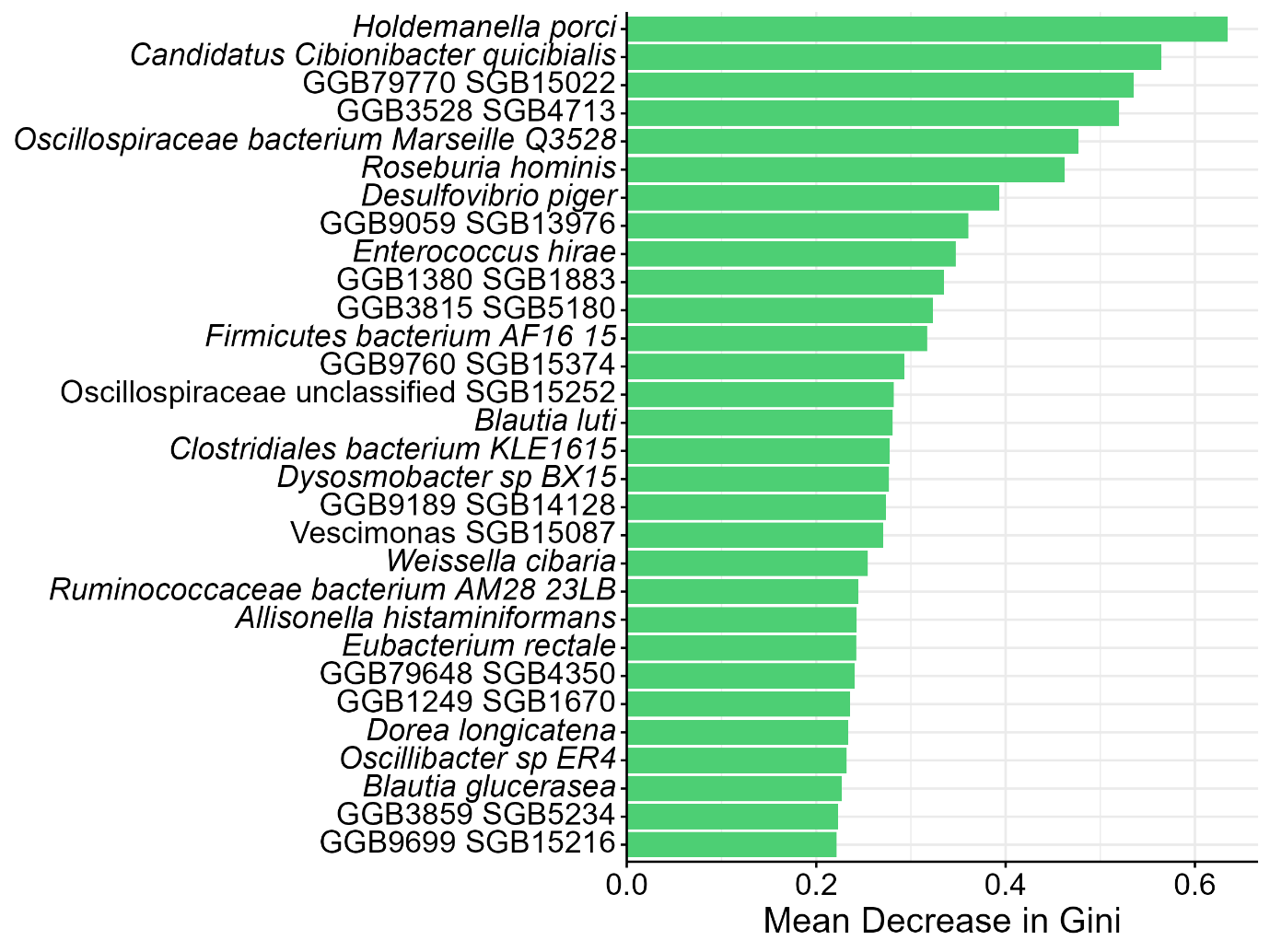

**Supplementary Figure 8:** Level of importance of bacterial species in separation of the BL and FU-associated taxonomic profiles in the ALB arm by a Random Forest-based approach represented by the mean decrease in Gini. The species are listed in order of importance from top to bottom.

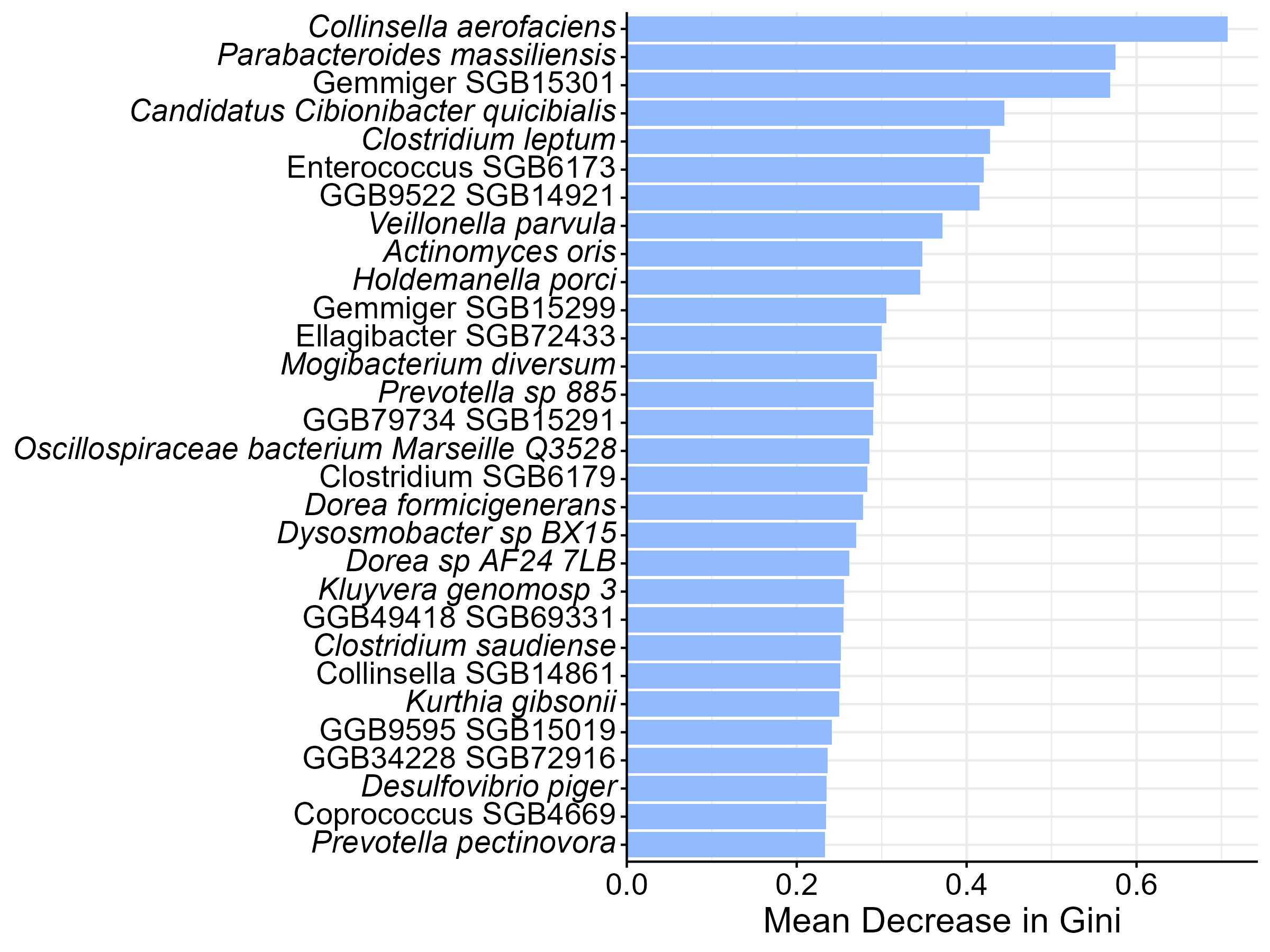

**Supplementary Figure 9:** Level of importance of bacterial species in separation of the BL and FU-associated taxonomic profiles in the IVM-ALB arm by a Random Forest-based approach represented by the mean decrease in Gini. The species are listed in order of importance from top to bottom.

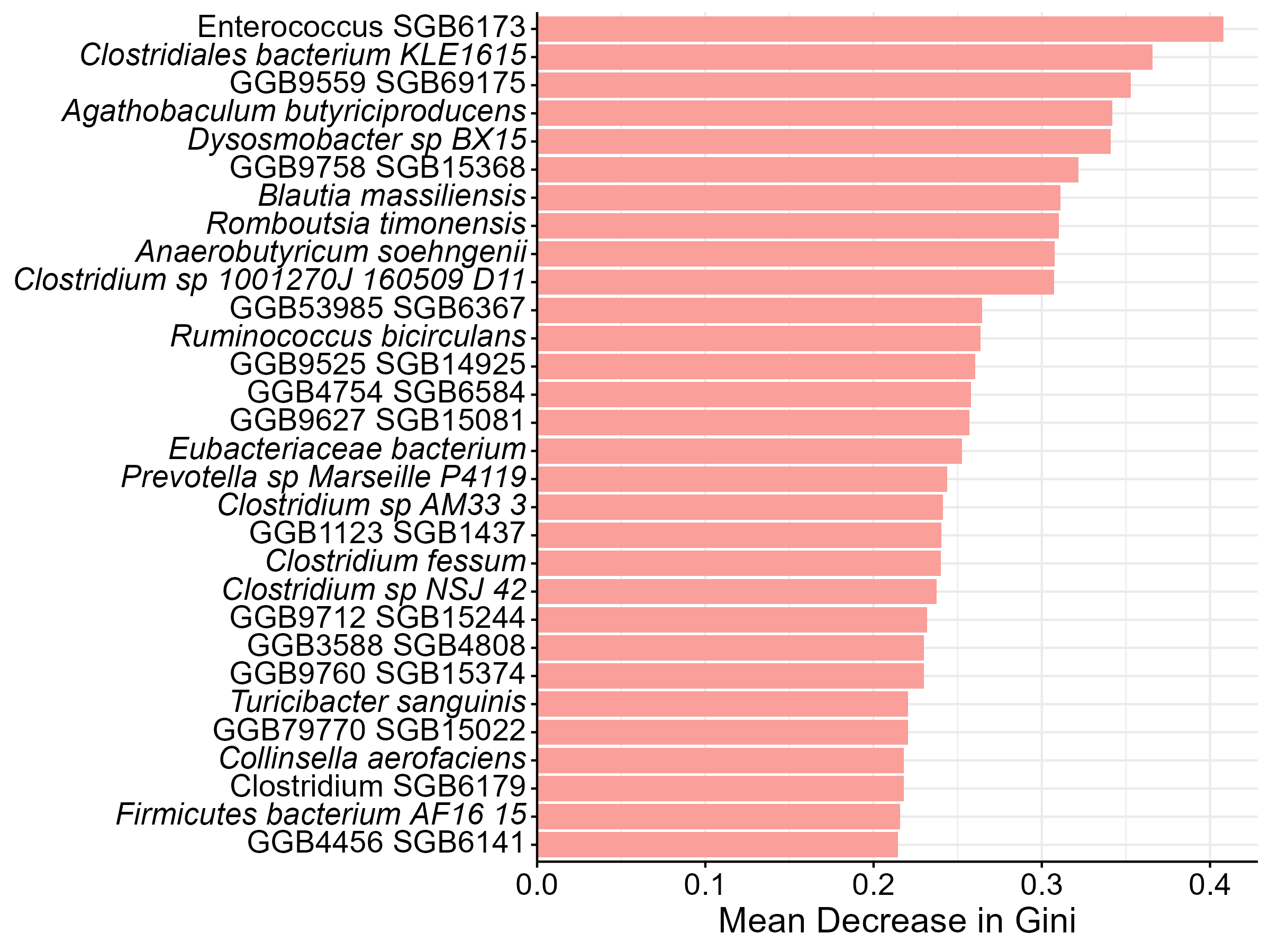

**Supplementary Figure 10:** Level of importance of bacterial species in separation of the BL and FU-associated taxonomic profiles in the MOX-ALB arm by a Random Forest-based approach represented by the mean decrease in Gini. The species are listed in order of importance from top to bottom.

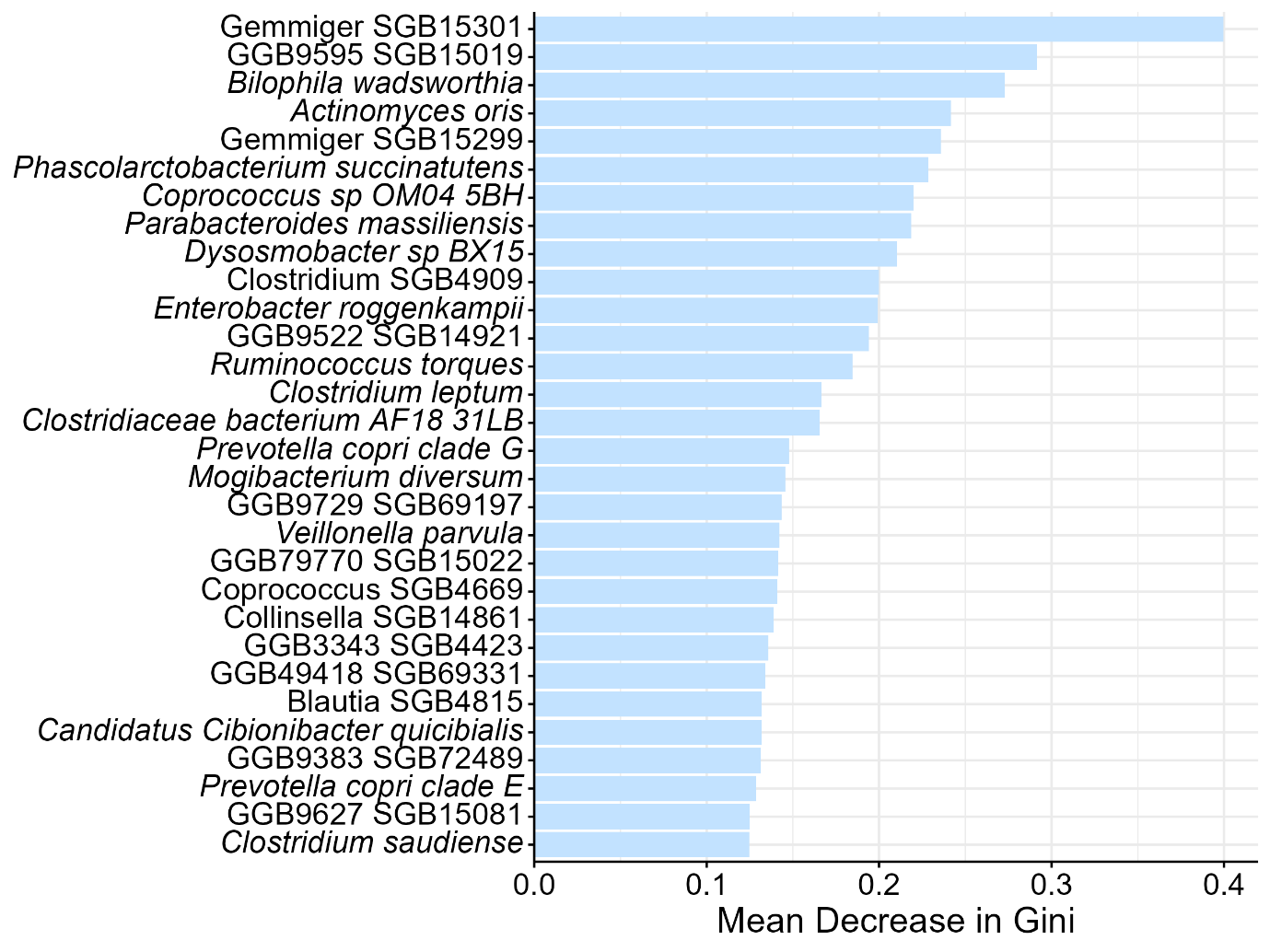

**Supplementary Figure 11:** Level of importance of bacterial species in separation of the BL and FU-associated taxonomic profiles in the “low” dosage category of the IVM-ALB arm by a Random Forest-based approach represented by the mean decrease in Gini. The species are listed in order of importance from top to bottom.

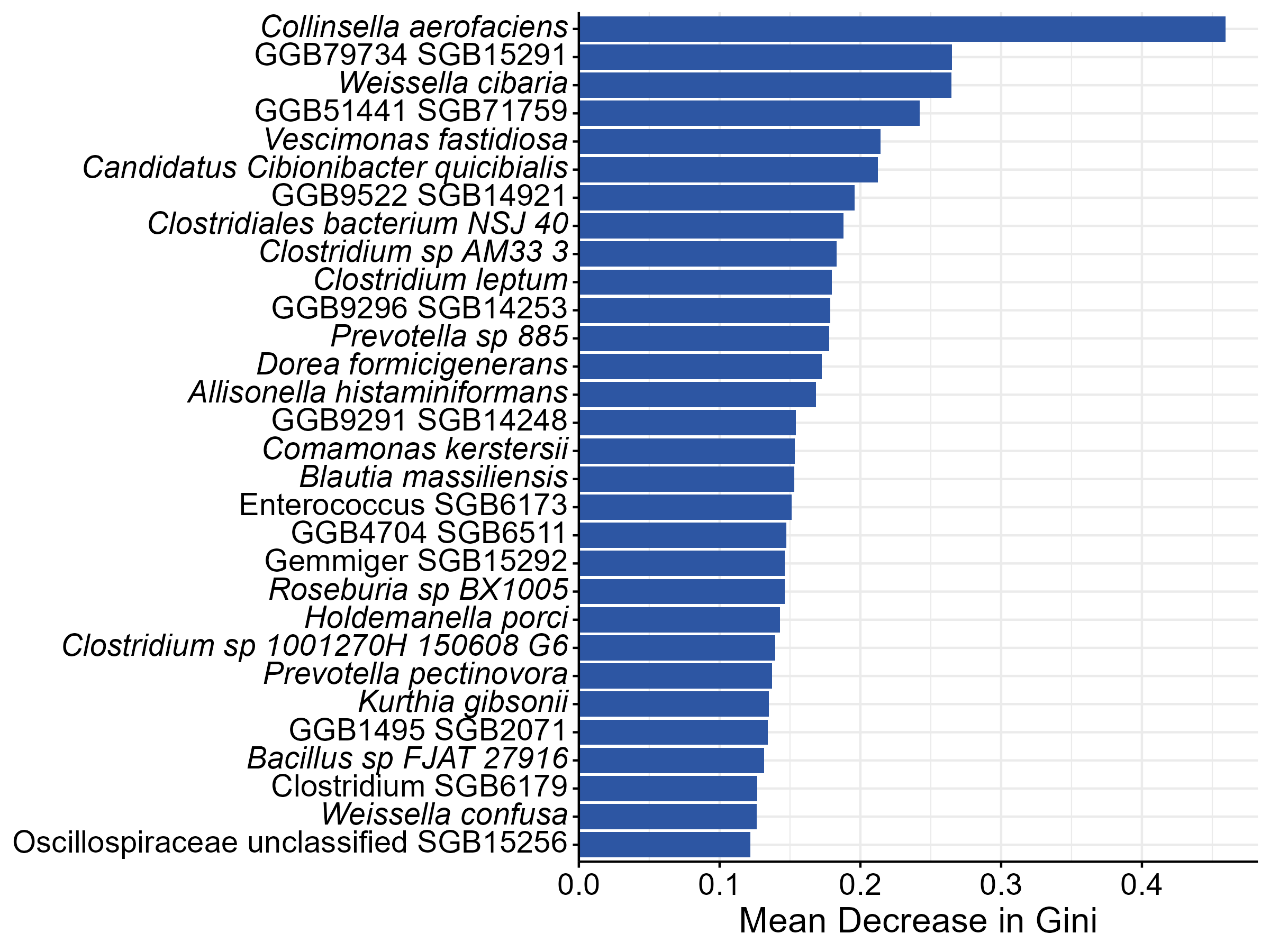

**Supplementary Figure 12:** Level of importance of bacterial species in separation of the BL and FU-associated taxonomic profiles in the “high” dosage category of the IVM-ALB arm by a Random Forest-based approach represented by the mean decrease in Gini. The species are listed in order of importance from top to bottom.

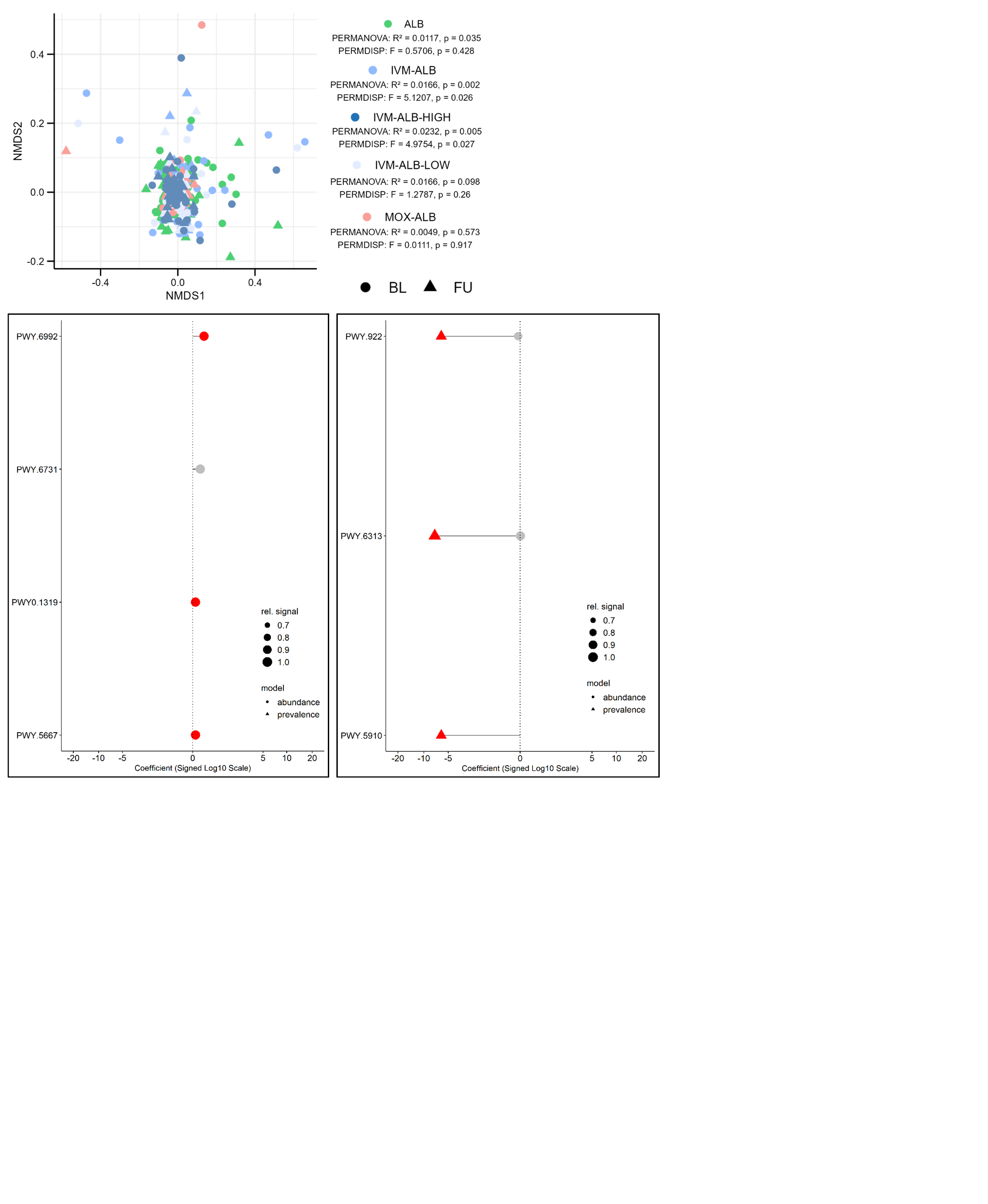

**Supplementary Figure 13:** Non-metric multidimensional scaling (NMDS) ordination based on Bray-Curtis Dissimilarity matrices of relative pathway abundances of paired baseline (BL) and follow-up (FU) samples. The stratification of IVM-ALB into IVM-ALB-LOW and IVM-ALB-HIGH is based on a weight threshold of 60kg, corresponding to 12mg of IVM. PERMANOVA: p = 0.035 (ALB), 0.002 (IVM-ALB), 0.005 (IVM-ALB-HIGH), 0.098 (IVM-ALB-LOW), 0.573 (MOX-ALB). PERMDISP: p = 0.428 (ALB), 0.026 (IVM-ALB), 0.027 (IVM-ALB-HIGH), 0.260 (IVM-ALB-LOW), 0.917 (MOX-ALB).

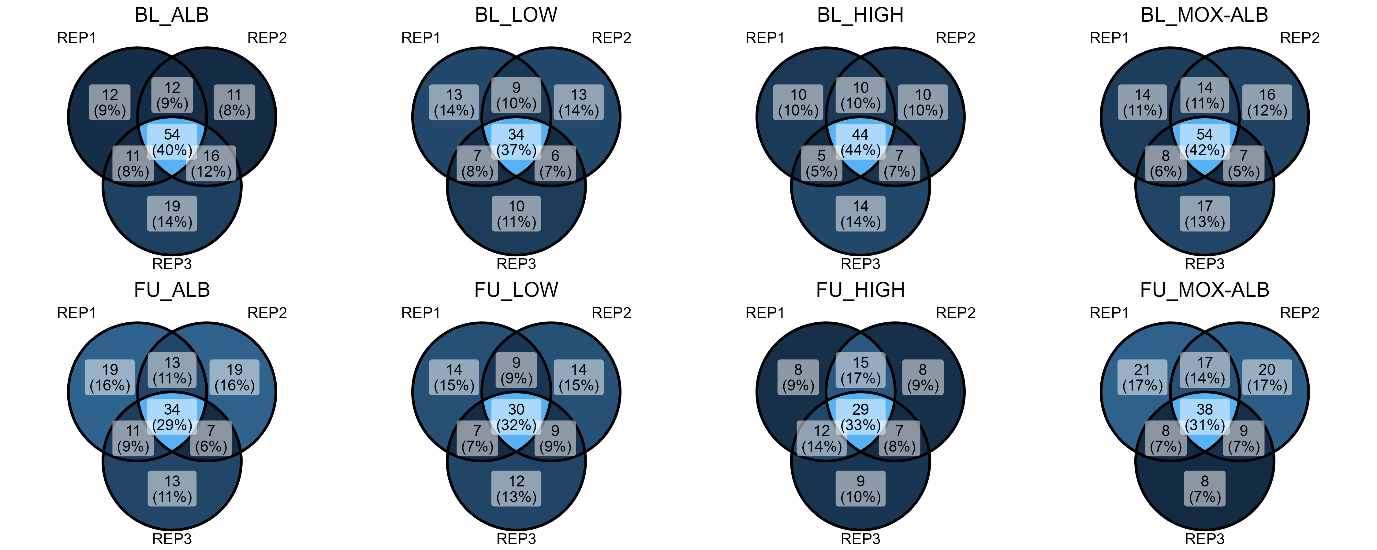

**Supplementary Figure 14:** Absolute number of shared and unique metagenomic bins for each replicate of a treatment arm (ALB, IVM-ALB-LOW, IVM-ALB-HIGH, MOX-ALB) and timepoint (BL, FU).

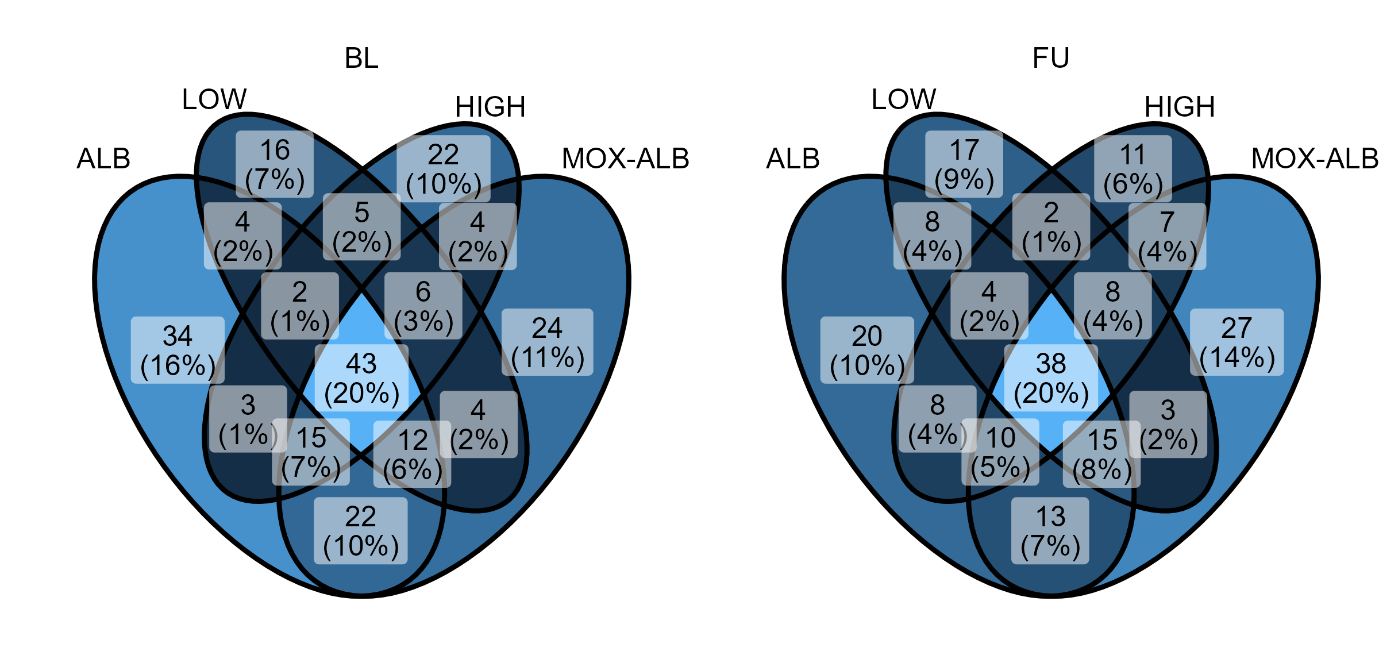

**Supplementary Figure 15:** Absolute number of shared and unique metagenomic bins between treatment arms (ALB, IVM-ALB-LOW, IVM-ALB-HIGH, MOX-ALB).

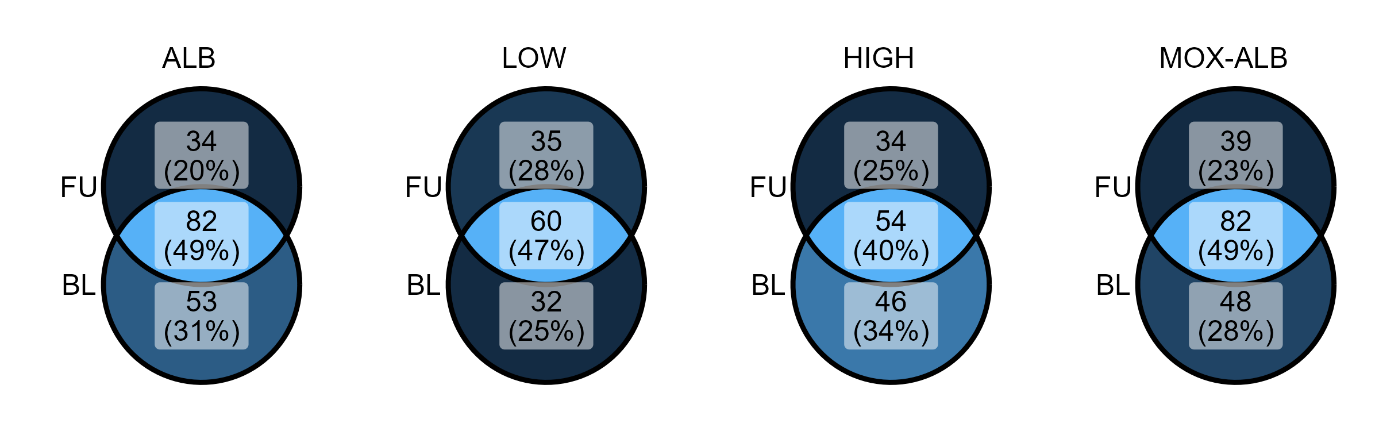

**Supplementary Figure 16:** Absolute number of shared and unique metagenomic bins between timepoints (BL, FU).

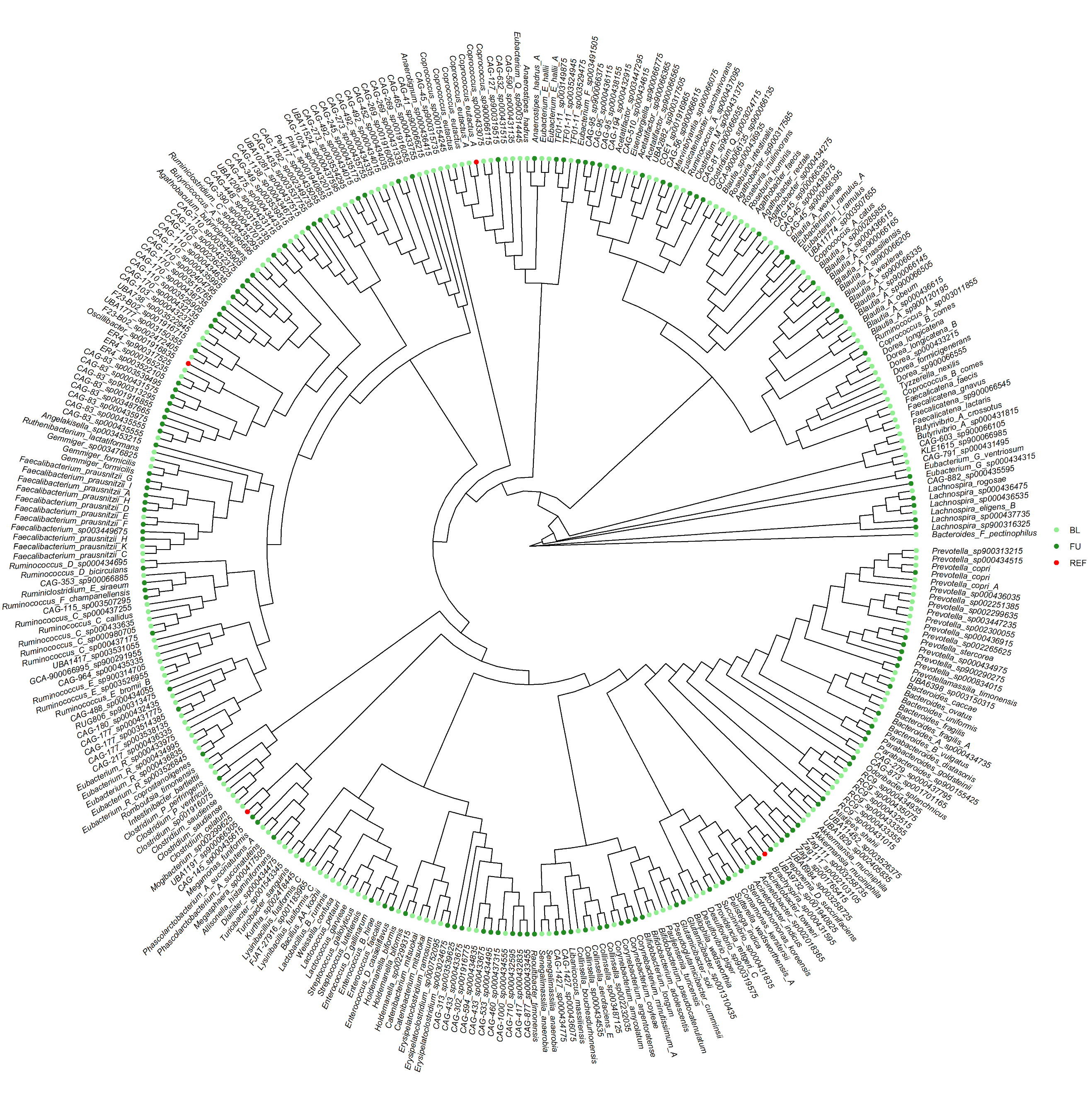

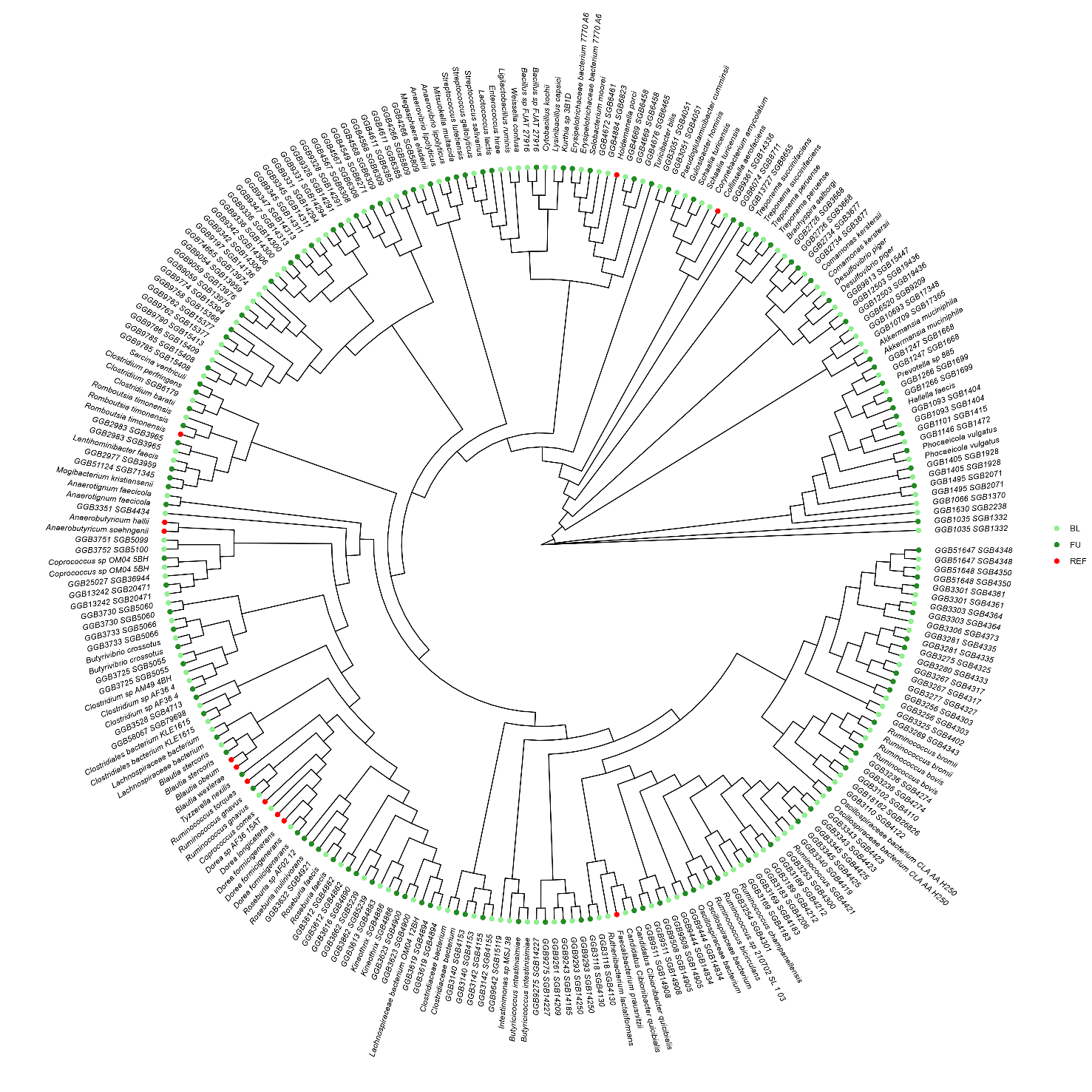

**Supplementary Figure 17:** Phylogenetic tree of the clean metagenomic assembled genomes (MAGs; Completeness > 95%, Contamination < 5%) of the ALB cohort. MAGs are colored based on the timepoint (BL = baseline, FU = follow-up). Bootstrapping values (n = 1000) are indicated at the tree nodes. Reference genomes of the identified core species were obtained via the CHOCOPhlAn database (Jan 2021) and are colored in red.

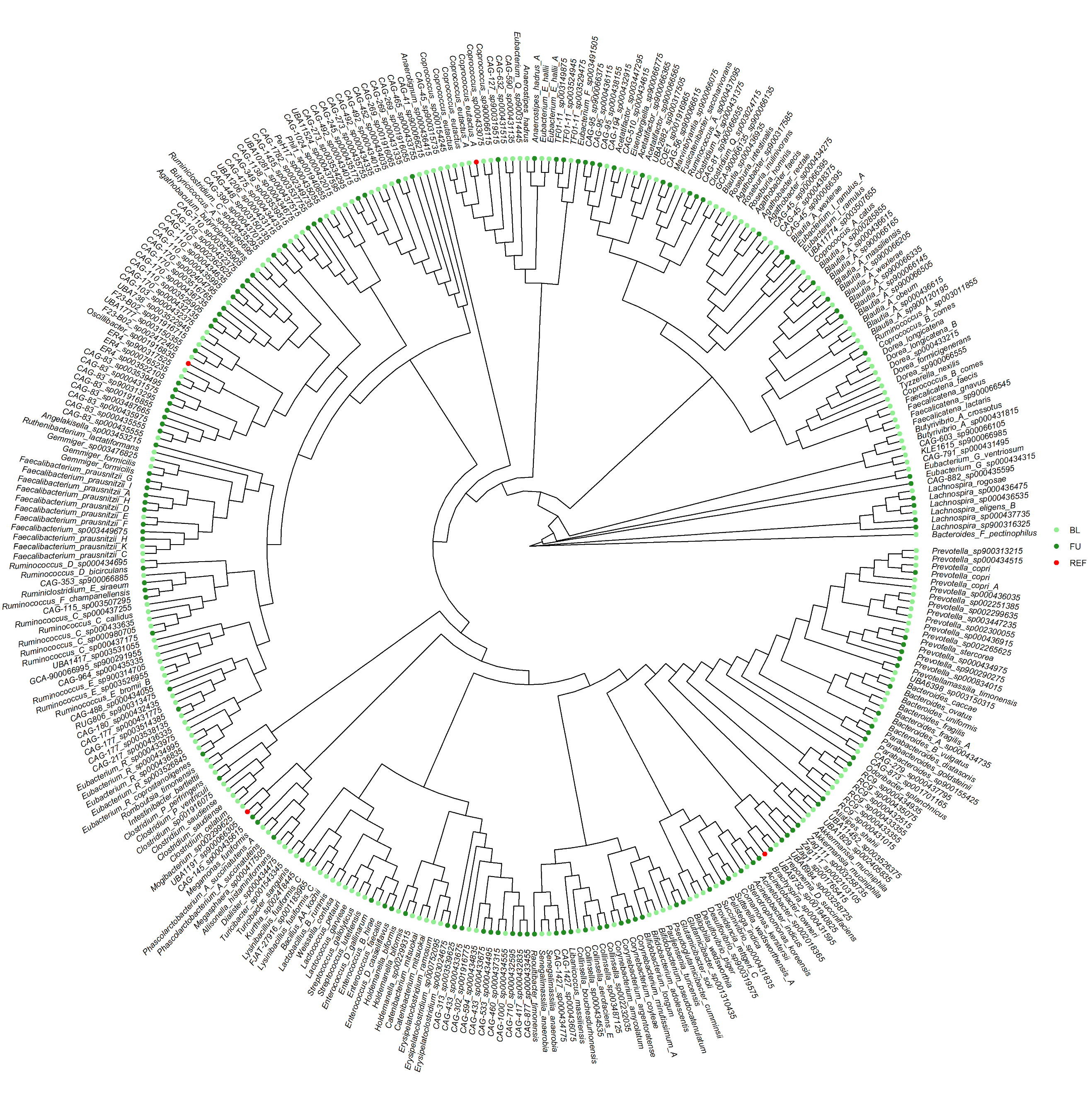

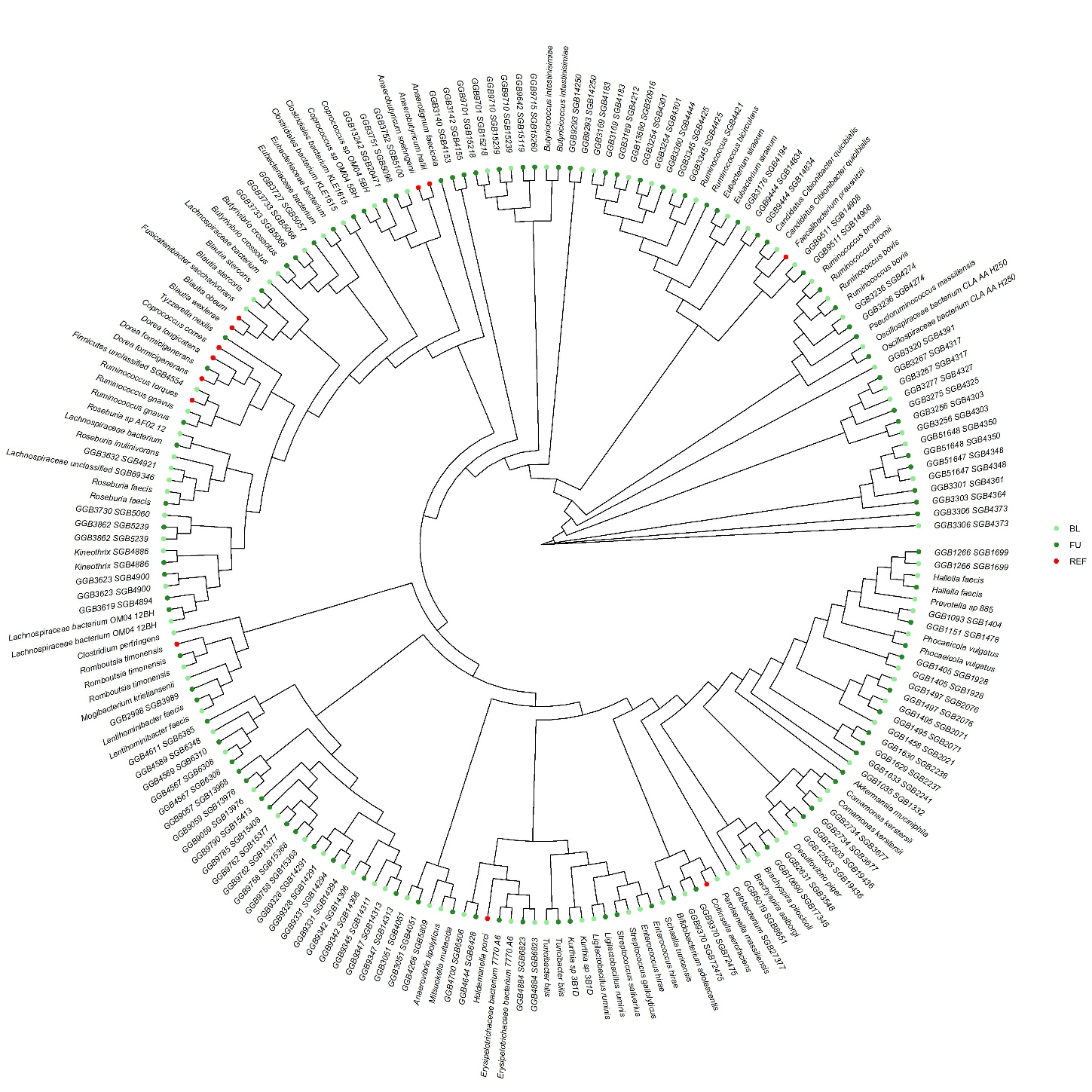

**Supplementary Figure 18:** Phylogenetic tree of the clean metagenomic assembled genomes (MAGs; Completeness > 95%, Contamination < 5%) of the IVM-ALB-LOW cohort. MAGs are colored based on the timepoint (BL = baseline, FU = follow-up). Bootstrapping values (n = 1000) are indicated at the tree nodes. Reference genomes of the identified core species were obtained via the CHOCOPhlAn database (Jan 2021) and are colored in red.

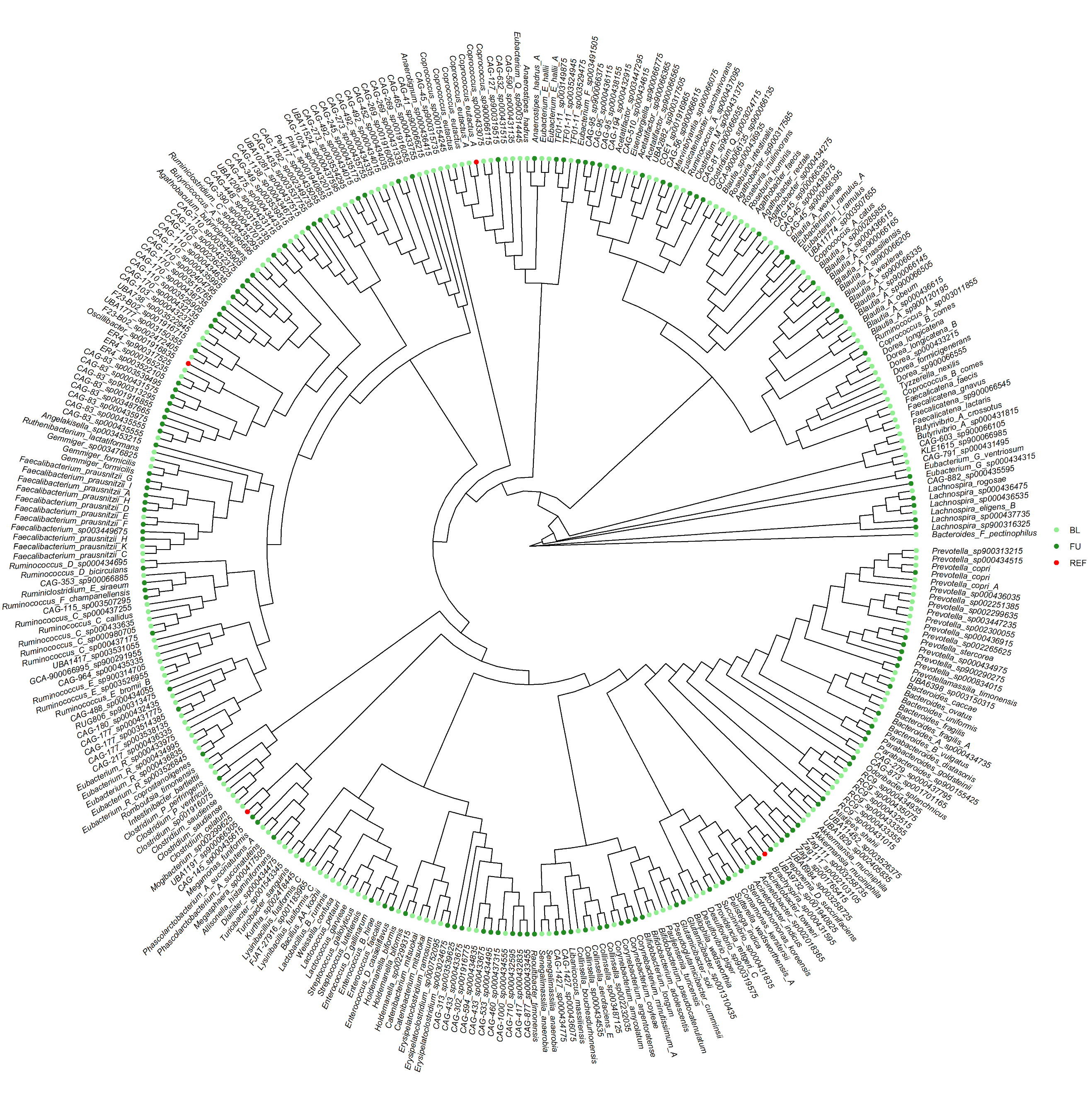

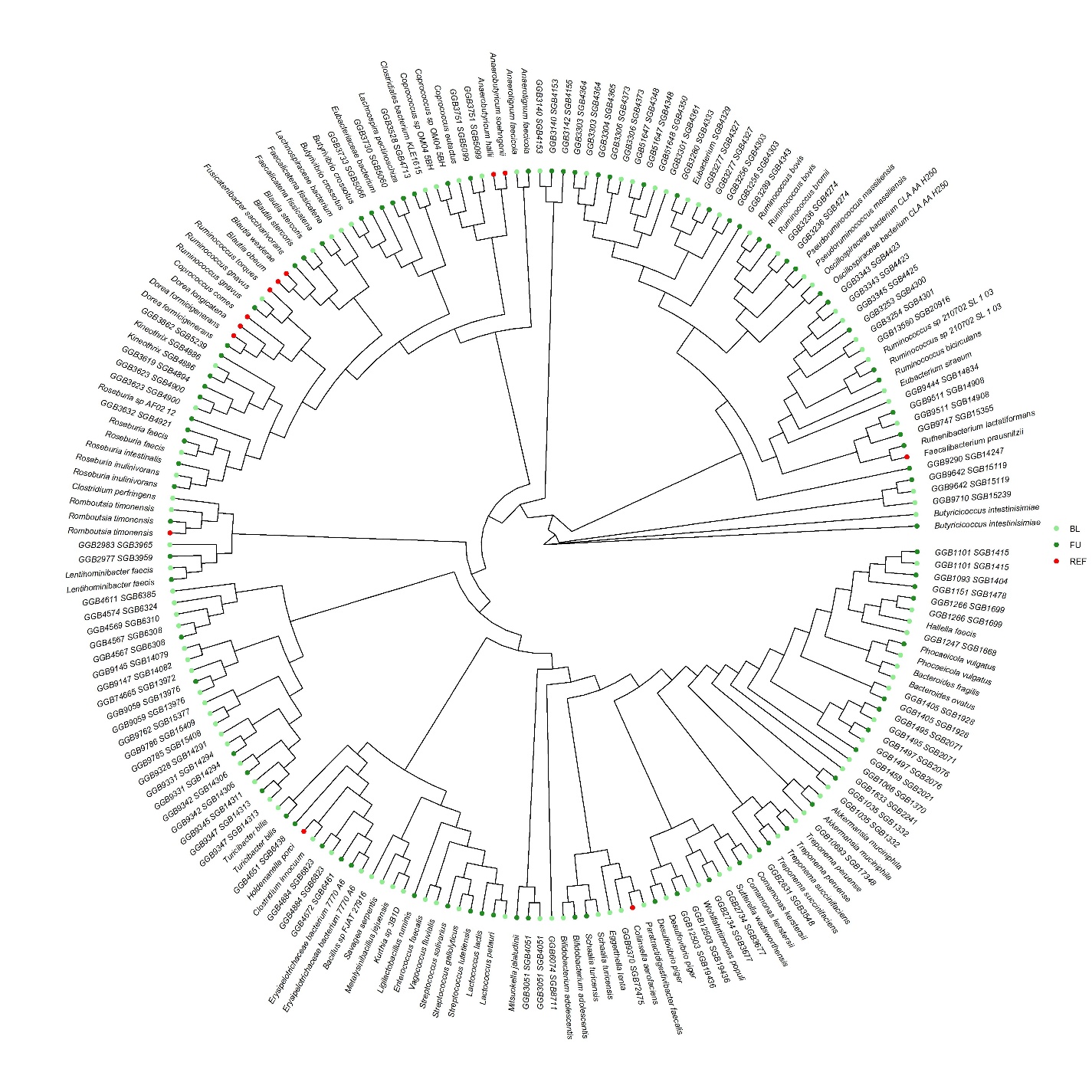

**Supplementary Figure 19:** Phylogenetic tree of the clean metagenomic assembled genomes (MAGs; Completeness > 95%, Contamination < 5%) of the IVM-ALB-HIGH cohort. MAGs are colored based on the timepoint (BL = baseline, FU = follow-up). Bootstrapping values (n = 1000) are indicated at the tree nodes. Reference genomes of the identified core species were obtained via the CHOCOPhlAn database (Jan 2021) and are colored in red.

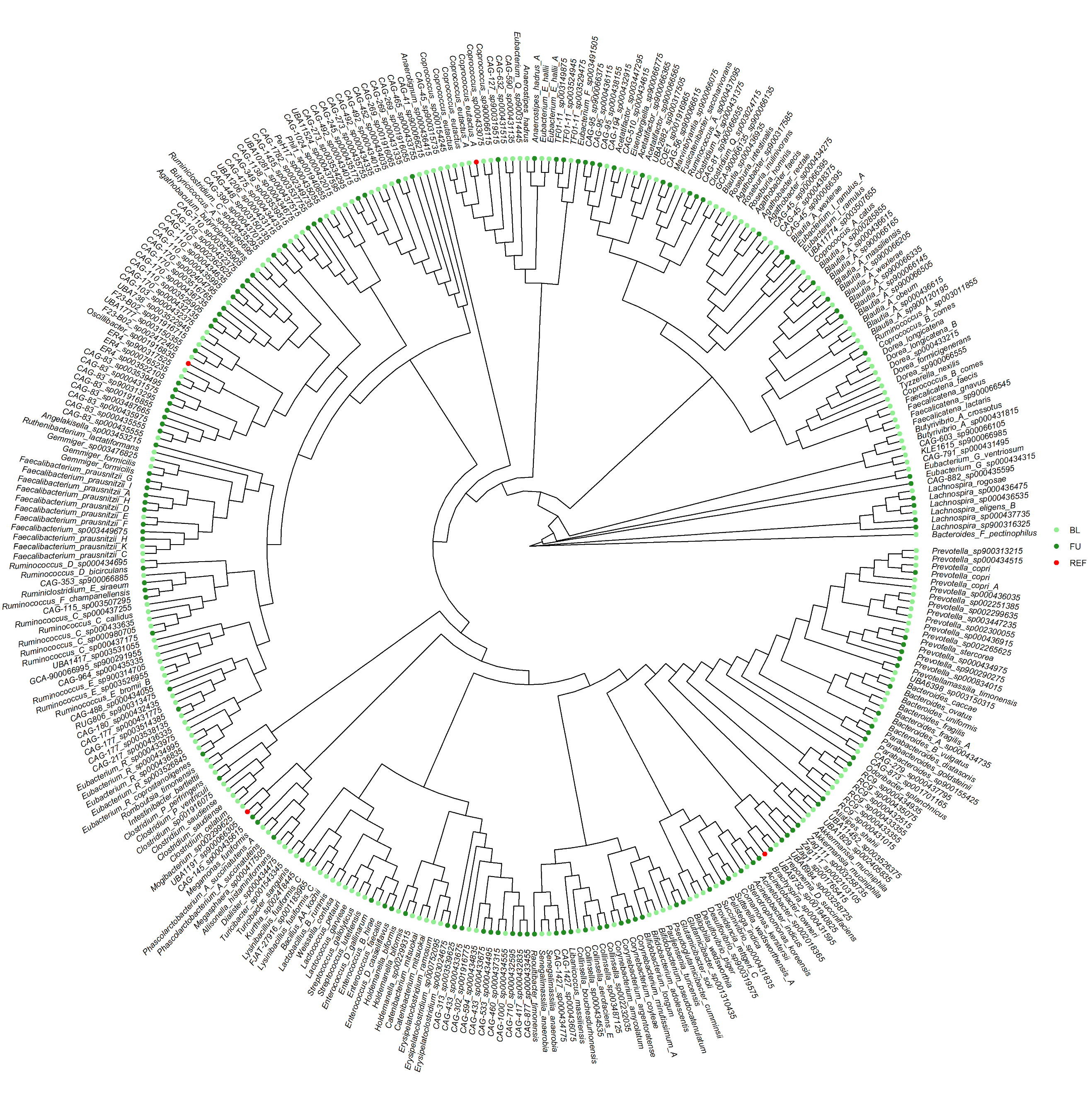

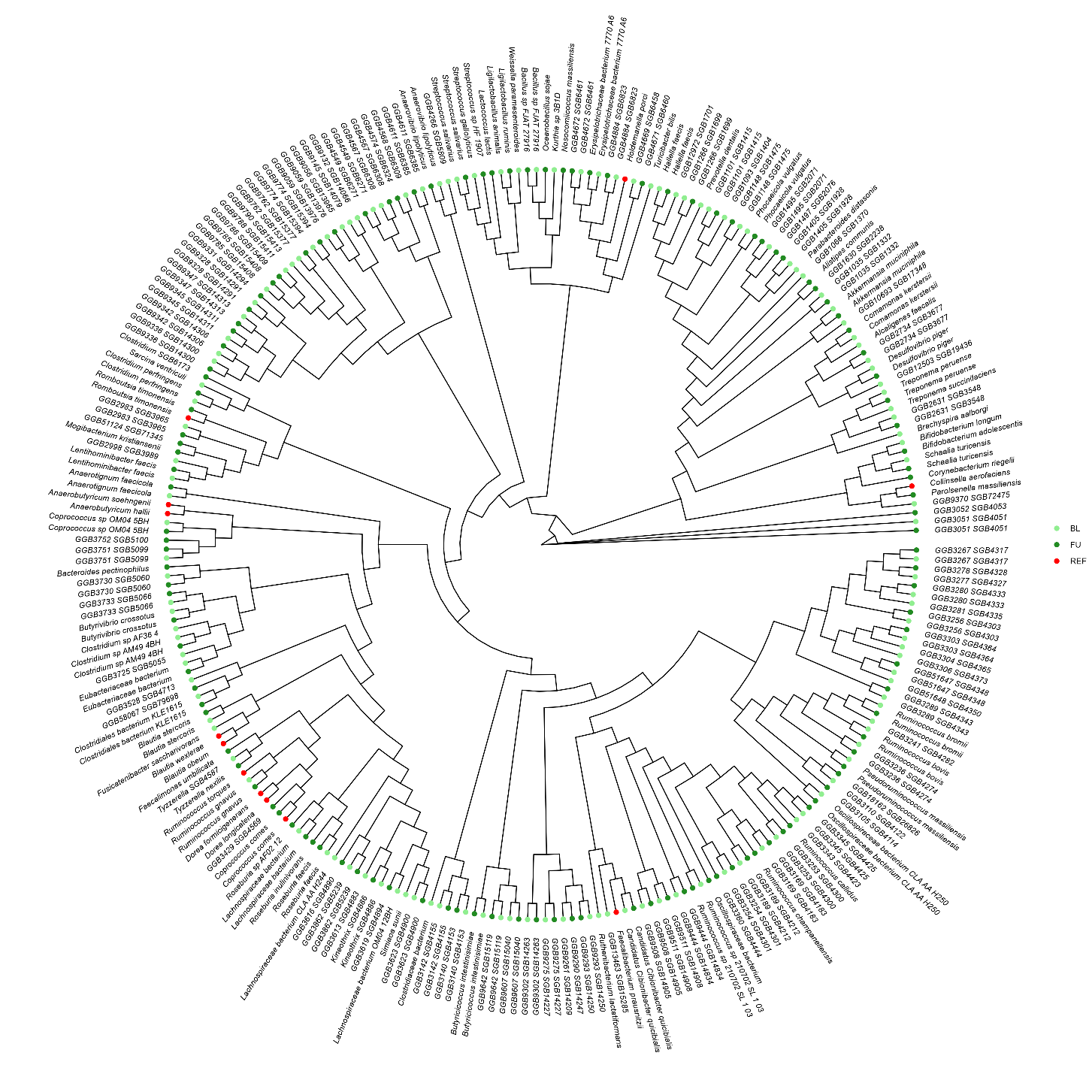

**Supplementary Figure 20:** Phylogenetic tree of the clean metagenomic assembled genomes (MAGs; Completeness > 95%, Contamination < 5%) of the MOX-ALB cohorts. MAGs are colored based on the timepoint (BL = baseline, FU = follow-up). Bootstrapping values (n = 1000) are indicated at the tree nodes. Reference genomes of the identified core species were obtained via the CHOCOPhlAn database (Jan 2021) and are colored in red.

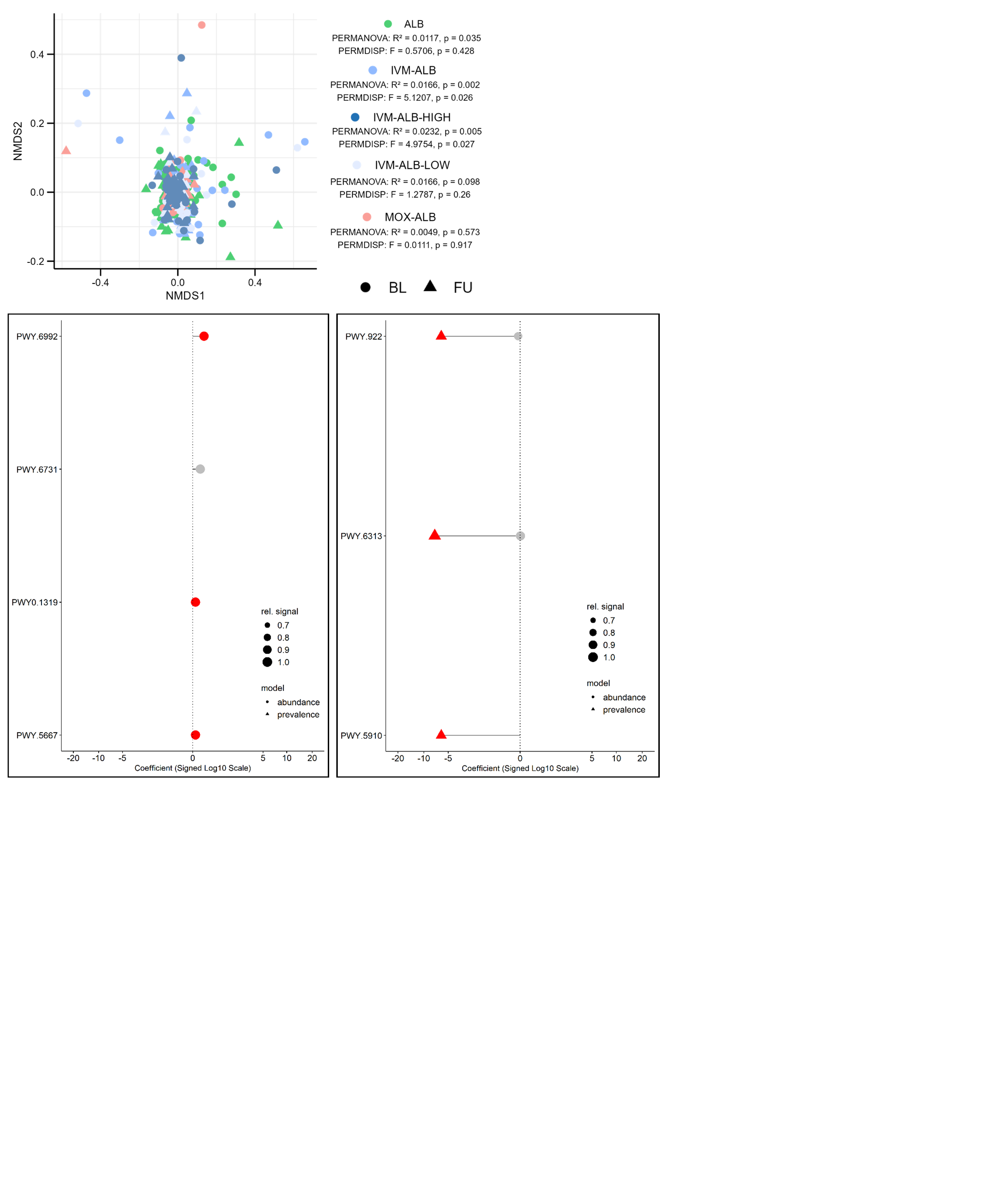

**Supplementary Figure 21:** Pairwise, species-level abundance and prevalence modelling of pathways between timepoints of ALB (left) and MOX-ALB (right) after filtering (keeping only the associations with the 20% lowest and 20% highest MaAsLin3 coefficients and a relative signal > 0.5). The relative signal was calculated as the ratio of samples, in which the association was detected divided by the total number of samples for that arm and timepoint. Only pathways reaching a “joint q-val” of < 0.1 are shown. Pathways with “individual q-val” < 0.1 are depicted in red. Coefficients for the abundance models are given as log2(FU/BL). Coefficients for the prevalence models are given as ln(OR). D. ALB E. IVM-ALB-HIGH F. MOX-ALB. IVM-ALB-LOW did not produce any significant results.

**Supplementary Figure 22:** Enriched or depleted MetaCyc pathways put forth by MaAsLin3 for each treatment arm. **Right:** Same enriched or depleted pathways after filtering, keeping only the associations with the 20% lowest and 80% highest MaAsLin3 coefficients for both abundance and prevalence modelling.

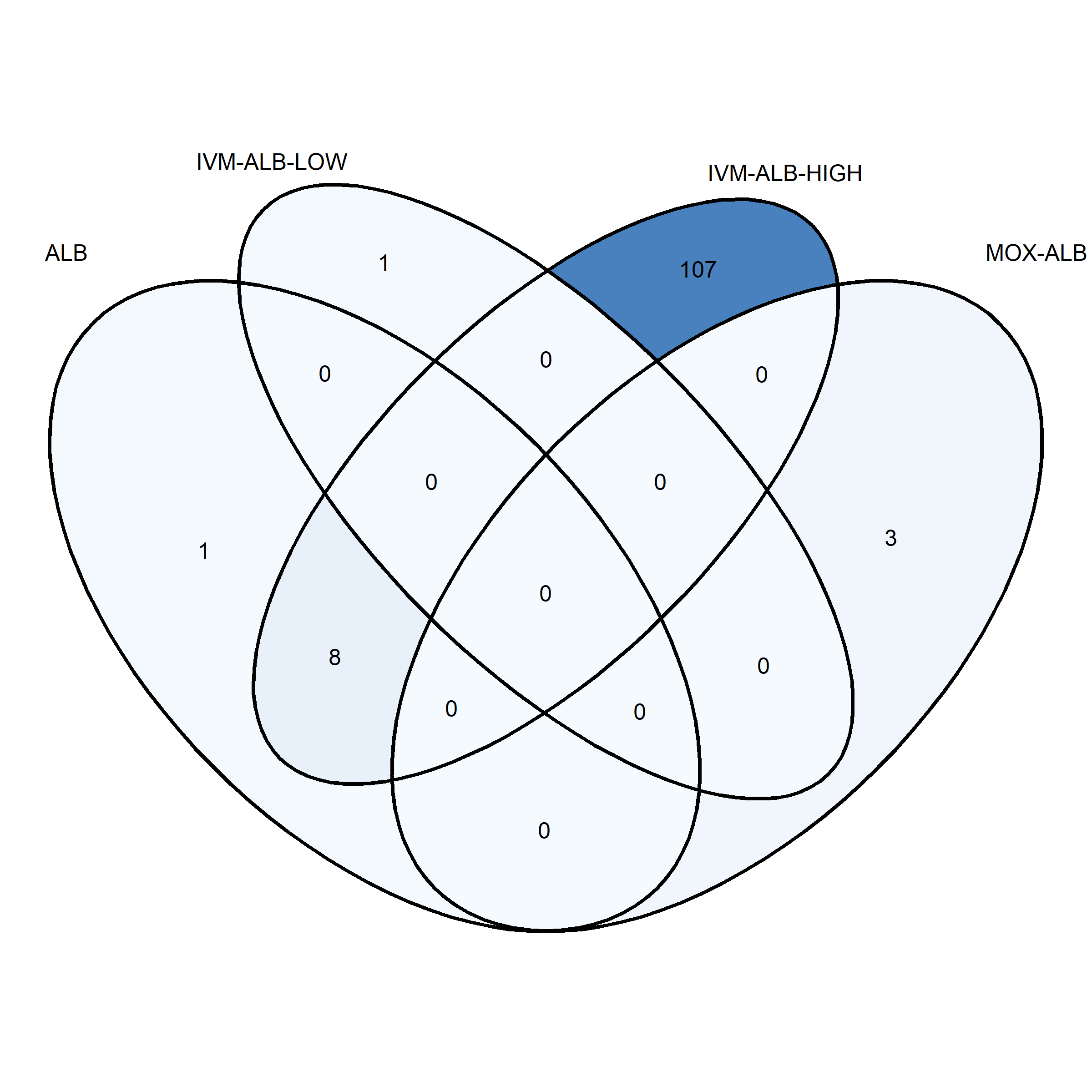

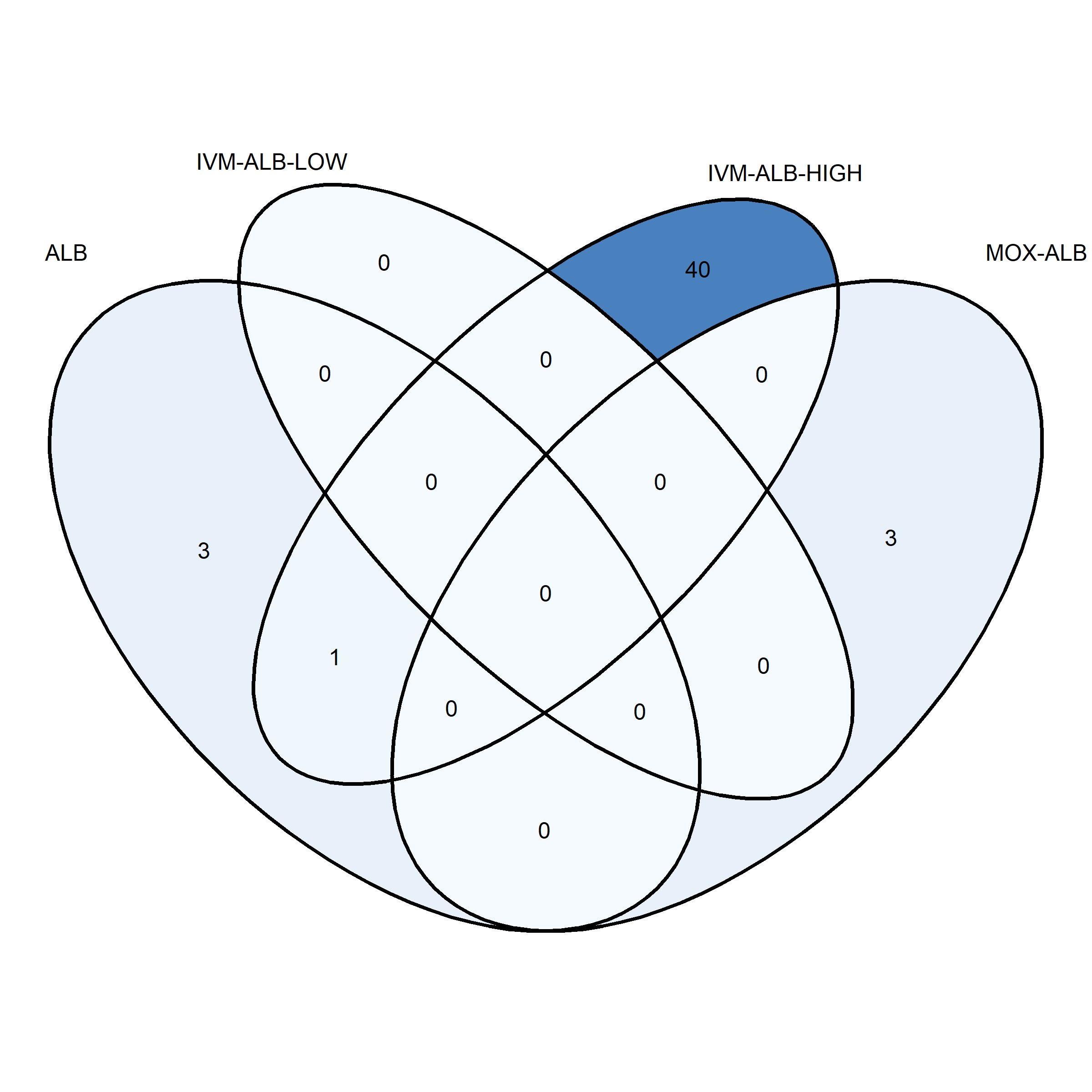

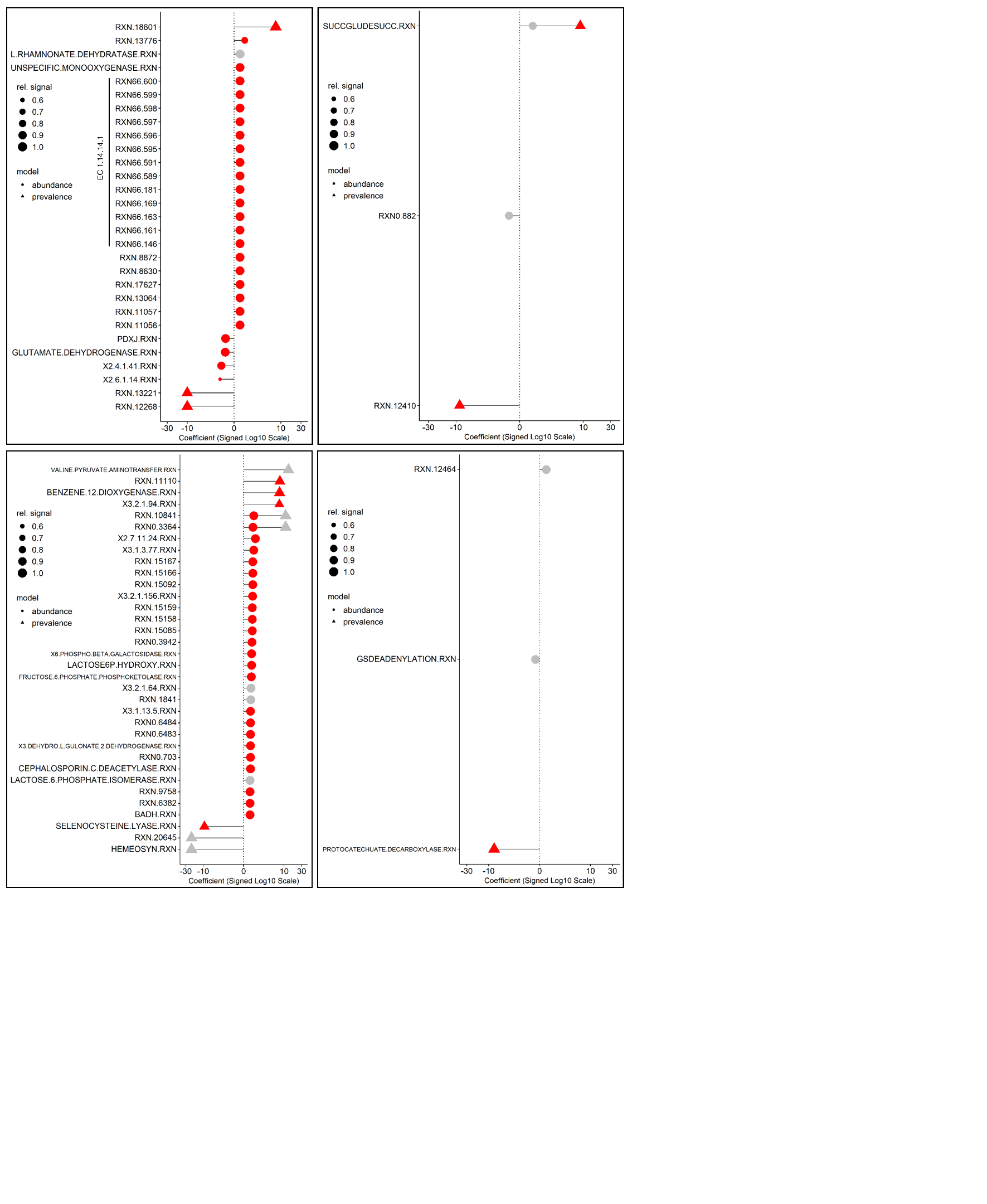

**Supplementary Figure 23:** Pairwise abundance and prevalence modelling of MetaCyc reactions between timepoints after filtering, keeping only the associations with the 5% lowest and 95% highest MaAsLin3 coefficients for both abundance and prevalence modelling and a relative signal > 0.5. The relative signal was calculated as the ratio of samples, in which the association was detected divided by the total number of samples for that arm and timepoint. Only reactions reaching a “joint q-val” of < 0.1 are shown. Reactions with “individual q-val” < 0.1 are depicted in red. Coefficients for the abundance models are given as log2(FU/BL). Coefficients for the prevalence models are given as ln(odds-ratio(FU/BL)). **Top-left to bottom-right:** ALB, IVM-ALB-LOW, IVM-ALB-HIGH, MOX-ALB.

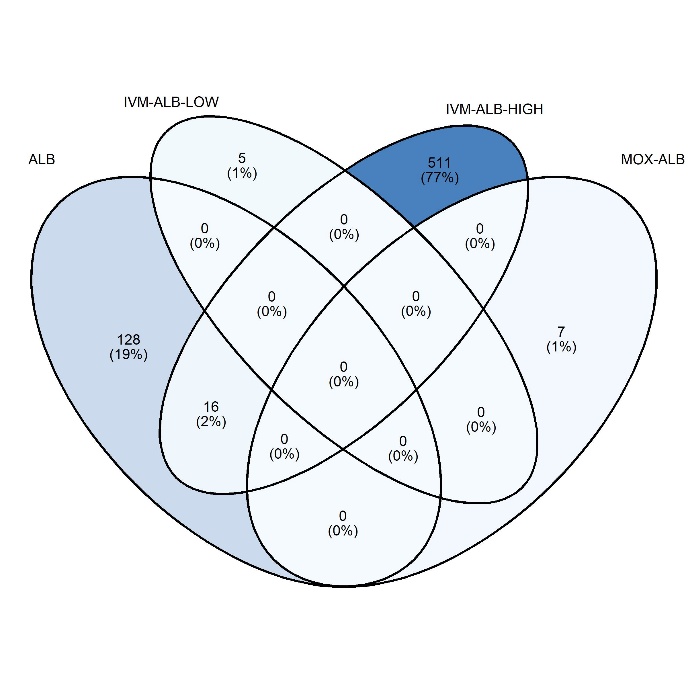

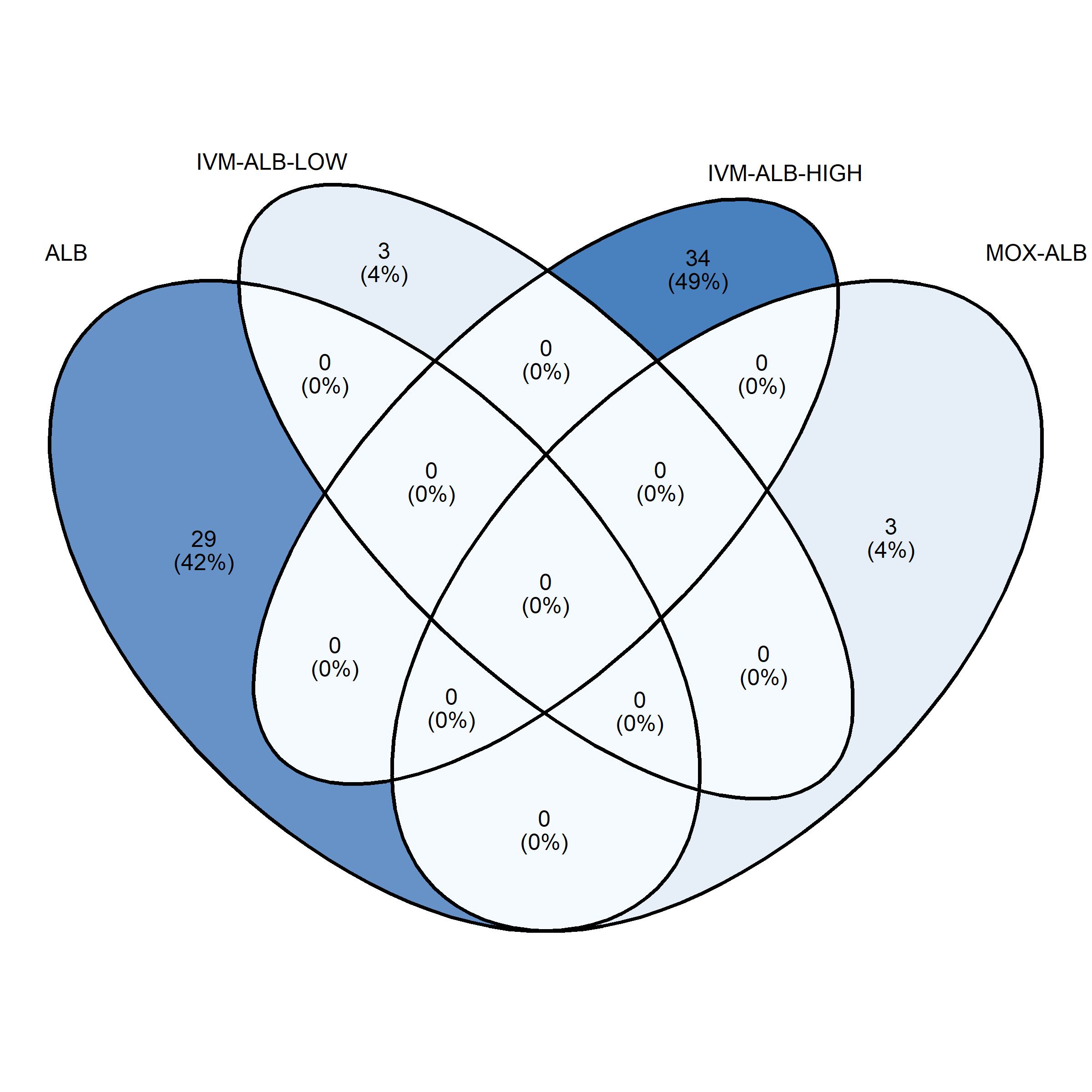

**Supplementary Figure 24: Left:** Enriched or depleted MetaCyc reactions put forth by MaAsLin3 for each treatment arm. **Right:** Same enriched or depleted reactions after filtering, keeping only the associations with the 5% lowest and 95% highest MaAsLin3 coefficients for both abundance and prevalence modelling.

**Supplementary Figure 25:** Non-metric multidimensional scaling (NMDS) ordination based on Bray-Curtis Dissimilarity matrices of paired BL and FU ARG abundances. The stratification of IVM-ALB into IVM-ALB-LOW and IVM-ALB-HIGH is based on a weight threshold of 60kg, corresponding to 12mg of IVM.

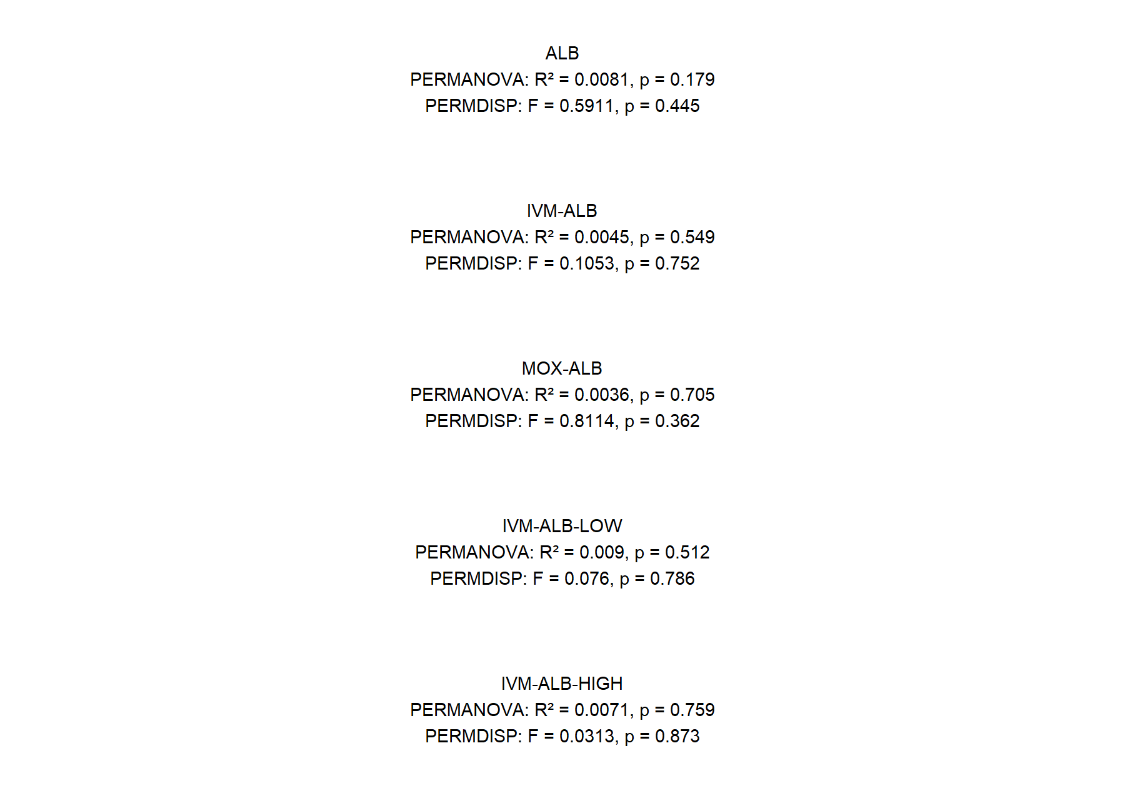

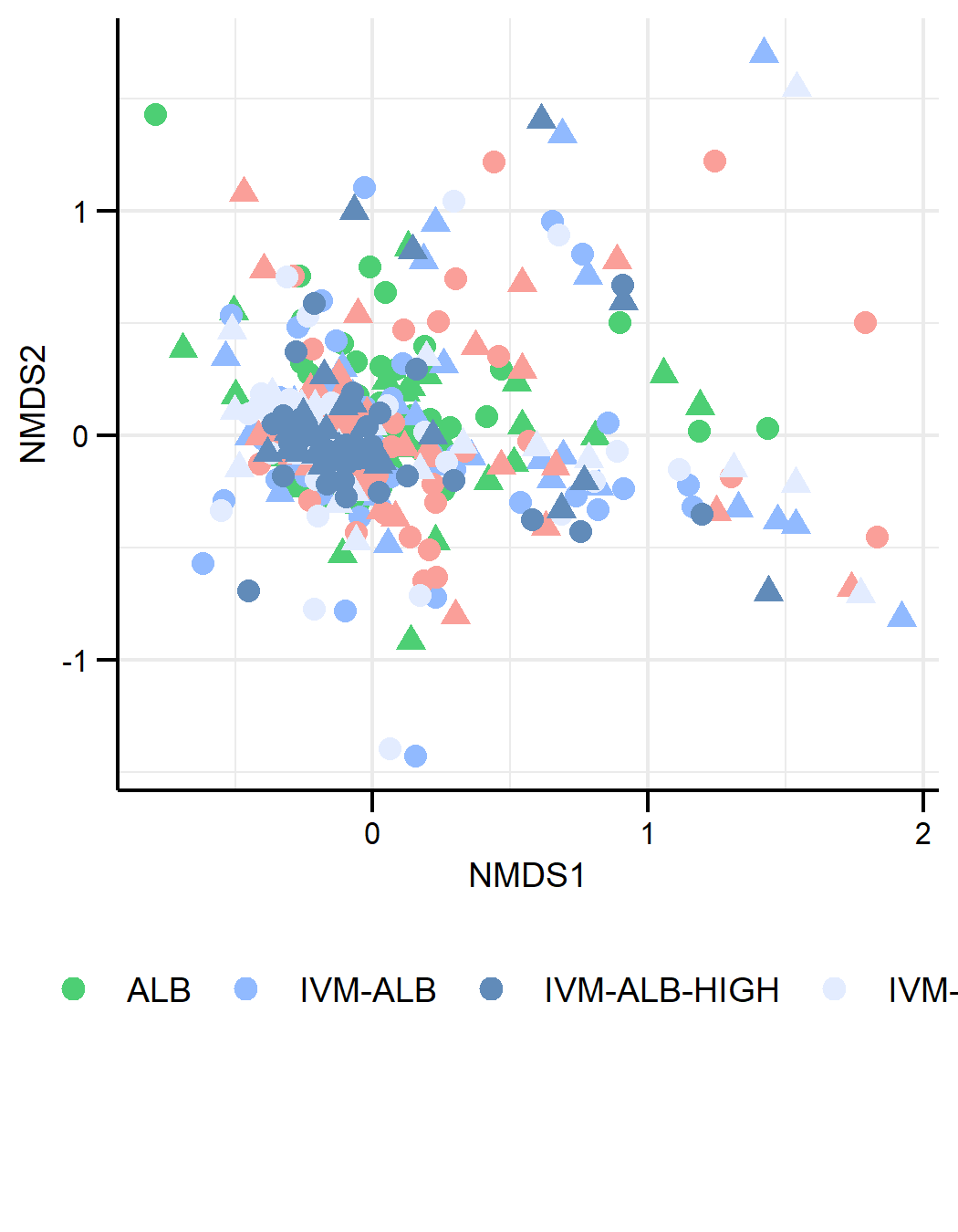

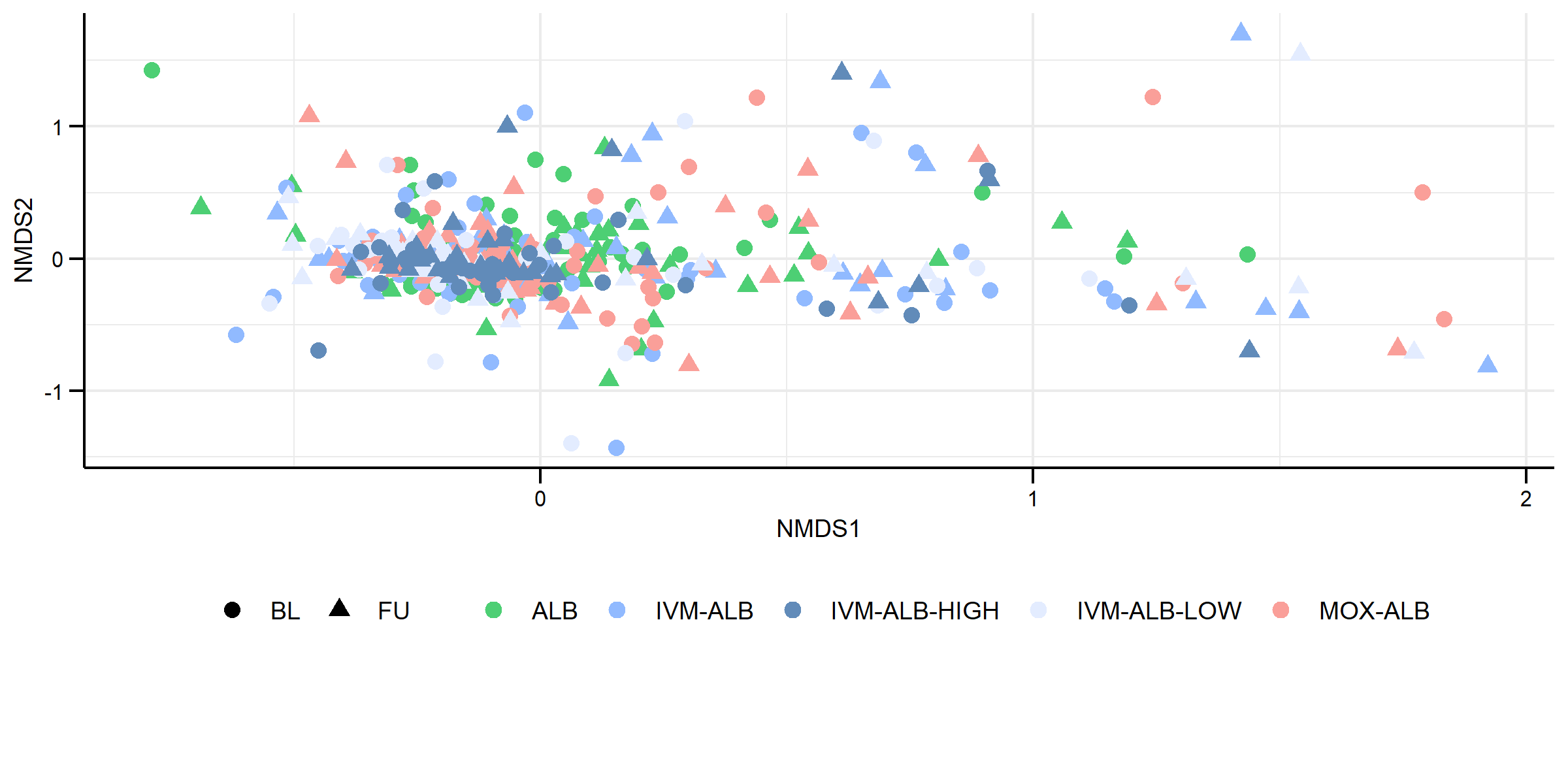
